# Growth regulation bringing modularity to morphogenesis of complex three-dimensional exoskeletons

**DOI:** 10.1101/2024.05.06.592547

**Authors:** Hiroshi C. Ito, Yu Uchiumi

## Abstract

Diverse three-dimensional morphologies of arthropods’ appendages, including beetle horns, are formed through the non-uniform growth of epidermis. Prior to moulting, epidermal tissue peels off from the old cuticle and grows non-uniformly to shape protruding appendages, which are often branching, curving, or twisting, from the planar epidermis. This non-uniform growth is possibly regulated by the distribution of morphogens on the epidermal cell sheet. Previous studies have identified molecules and signalling pathways related to such morphogenesis; however, how local regulation of cell sheet growth can transform planar epidermis globally into complex three-dimensional structures, such as beetle horns, remains unclear. To reveal the relationship between epidermal growth regulation and generated structures, this study theoretically examined how various shapes can be generated from planar epidermis under a deductive growth model that corresponds morphogen distributions to non-uniform growth on tissue. The results show that the heterochronic expression of multiple morphogens can flexibly fuse multiple simple shapes to generate various structures emulating complex appendages of beetles. These findings indicate that morphogenesis through such a mechanism may have developmental stability and modularity, providing insights into the evolution of the diverse morphology of arthropods.

## 1. Introduction

Arthropods have diverse three-dimensional (3D) morphologies. In particular, the morphology of their appendages varies among species, sexes, developmental stages, and environmental conditions, with beetle horns being a typical example [1]. Although horns often have extreme sizes and elaborate shapes, they can evolve rapidly and dramatically and are easily lost or gained evolutionarily [2]. Moreover, horns should have mechanical strength as a weapon of competition in males and developmental plasticity depending on the body size [3,4], implying that the morphogenesis has ambivalence between evolvability and developmental stability. Therefore, such appendages of arthropods are representative examples of a classical paradox in evolutionary developmental biology [5].

The morphogenesis of exoskeletal appendages, such as beetle horns, depends on the non-uniform growth of epidermal tissues during metamorphosis or moulting [6,7]. Prior to moulting, the epidermal cell sheet lining the old cuticle detaches from the cuticle and grows non-uniformly to transform from the planar shape constituting the larval epidermis into the shape of an adult appendage, producing a new cuticle [8]. Because the synthesised cuticle does not have strong bending elasticity, the growing cell sheet and new cuticle are compacted inside the old cuticle in a wrinkled and folded state [9]. They extend to form an elastic or rigid 3D shape during moulting, along with the break out of the old cuticle [9]. The non-uniform growth of epithelial sheets involves non-uniform cell division, rearrangement, and growth, which are integratively controlled to achieve harmonious 3D shapes [8]. Recent studies have identified genes and pathways involved in the integrative control to form the appendages [10–13]. For example, in the case of beetle horns [1], the genes related to leg patterning (e.g. *Distal-less* and *dachshund*) also contribute to the patterning of their proximo-distal axis; the Hox gene (*Sex-combs reduced*) affects their positioning; and several pathways, including insulin signalling, regulate their relative size. *Notch* and *CyclinE* affect the depth and pattern, respectively, of the primordial furrow of the epidermal tissue inside the prepupal cuticle, which is related to the proliferation of epidermal cells and determines the shape of the adult horn [14].

However, understanding how such non-uniform growth of epidermal sheets is controlled harmoniously to generate 3D structures, including horns and other appendages, remains challenging. Previous studies have shown how local cell mechanics allow the construction of 3D structures, e.g. the respiratory tubes of *Drosophila* eggs [15] and folding patterns similar to an insect primordium [16]. Another study revealed that the folding patterns on the primordium can expand into its horn structure [17]. Because these studies assume in advance certain distributions of mechanical forces, sheet growths, or folding patterns, how such distributions may be formed and regulated by morphogens have not been clarified. To reveal the morphogenesis of the appendages via non-uniform growth regulated by morphogen intensity distributions, Ito [18] proposed a ‘morphogen-gradient-squared (MGS) growth model’, in which the direction of sheet growth is parallel to the morphogen intensity gradient at that position and its growth rate is determined by the gradient strength. This model can transform a flat epidermal sheet into a 3D structure that materialises the shape of a morphogen distribution, if the shape is sufficiently simple such as a straight and unbranched horn or spine (figure 1). In addition, the generated shape can be further curved and twisted by incorporating the expression of additional morphogens (figure 2h, 2i). However, exoskeletal appendages of arthropods, such as beetle horns, have more complex shapes involving branches, and the morphogenesis of such appendages remains unclear, particularly in the relationship between growth regulation and resulting shape of an extended sheet.

**Figure 1.**
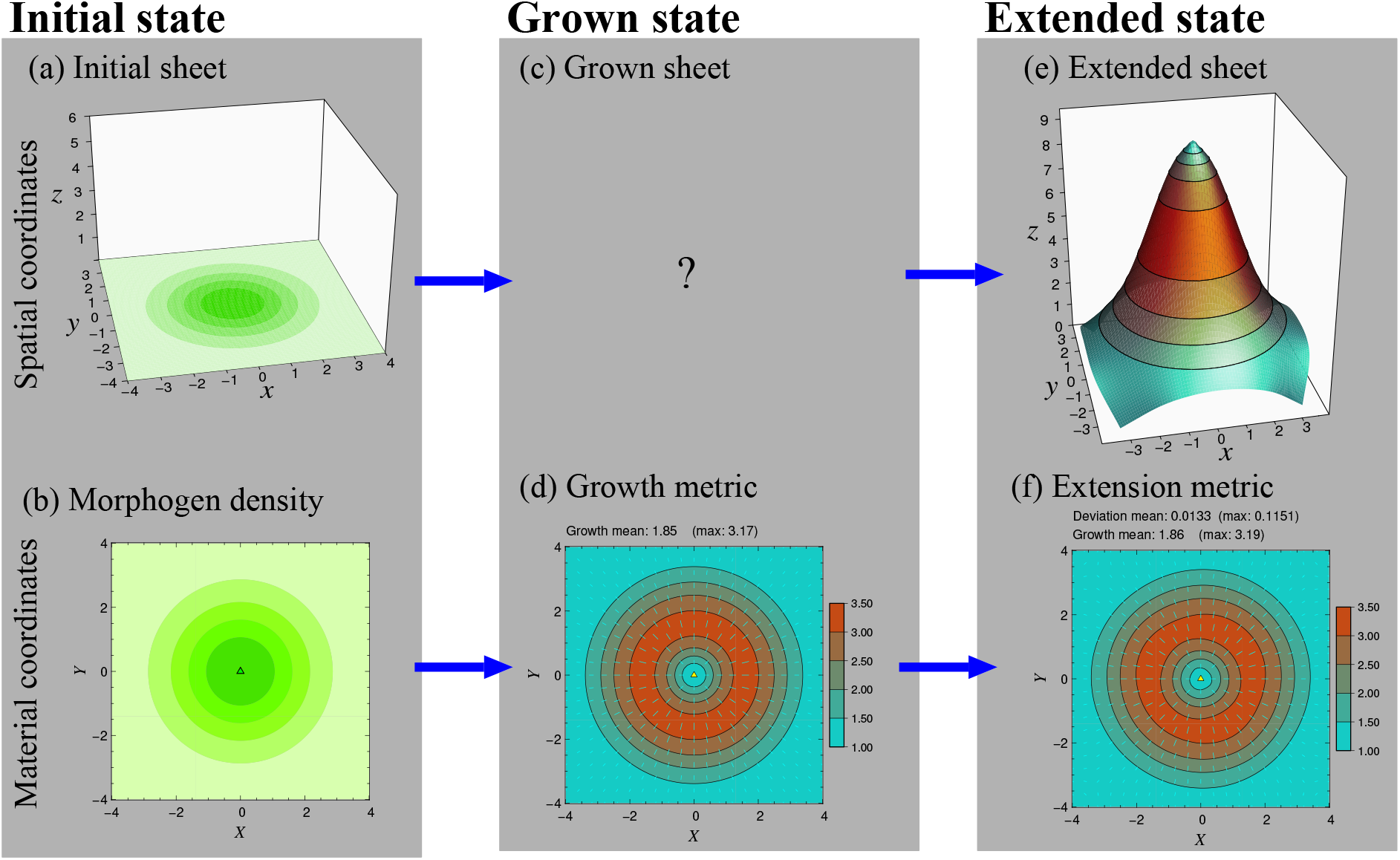
Initial state, grown state, and extended state of epidermal sheet in spatial coordinate system **r** = (*x,y,z*)^T^ and material coordinate system **u** = (*X,Y*)^T^. (a) Initial flat sheet in the spatial coordinates. Green gradient indicates morphogen intensity (light green: low, deep green: high). (b) Initial sheet in the material coordinates. Green gradient indicates morphogen intensity as in (a). The small triangle indicates the maximum morphogen intensity. (c) Grown sheet in the spatial coordinates (not analysed in this study and hence unknown). (d) Grown sheet in the material coordinates, represented with the growth metric **M**(**u**) given by equation (6). The cyan-red gradient indicates the square-root of the maximum eigenvalue of **M**(**u**) at each position (i.e., how many times each microelement of the sheet is elongated) with its eigenvector (i.e., growth direction) indicated with the short cyan line segment at that position. The small yellow triangle indicates a maxima of Riemann scalar curvature (expected to become a projection or follow in the extended sheet). (e) Extended sheet in the spatial coordinates. The cyan-red gradient indicates the maximum eigenvalue of **M**(**u**), and which is equivalent to (d). (f) Extended sheet in the material coordinates represented with the extension metric **M**_ext_(**u**) that was calculated from the extended sheet (Appendix B.1) and indicated in the same manner as in (d).

**Figure 2.**
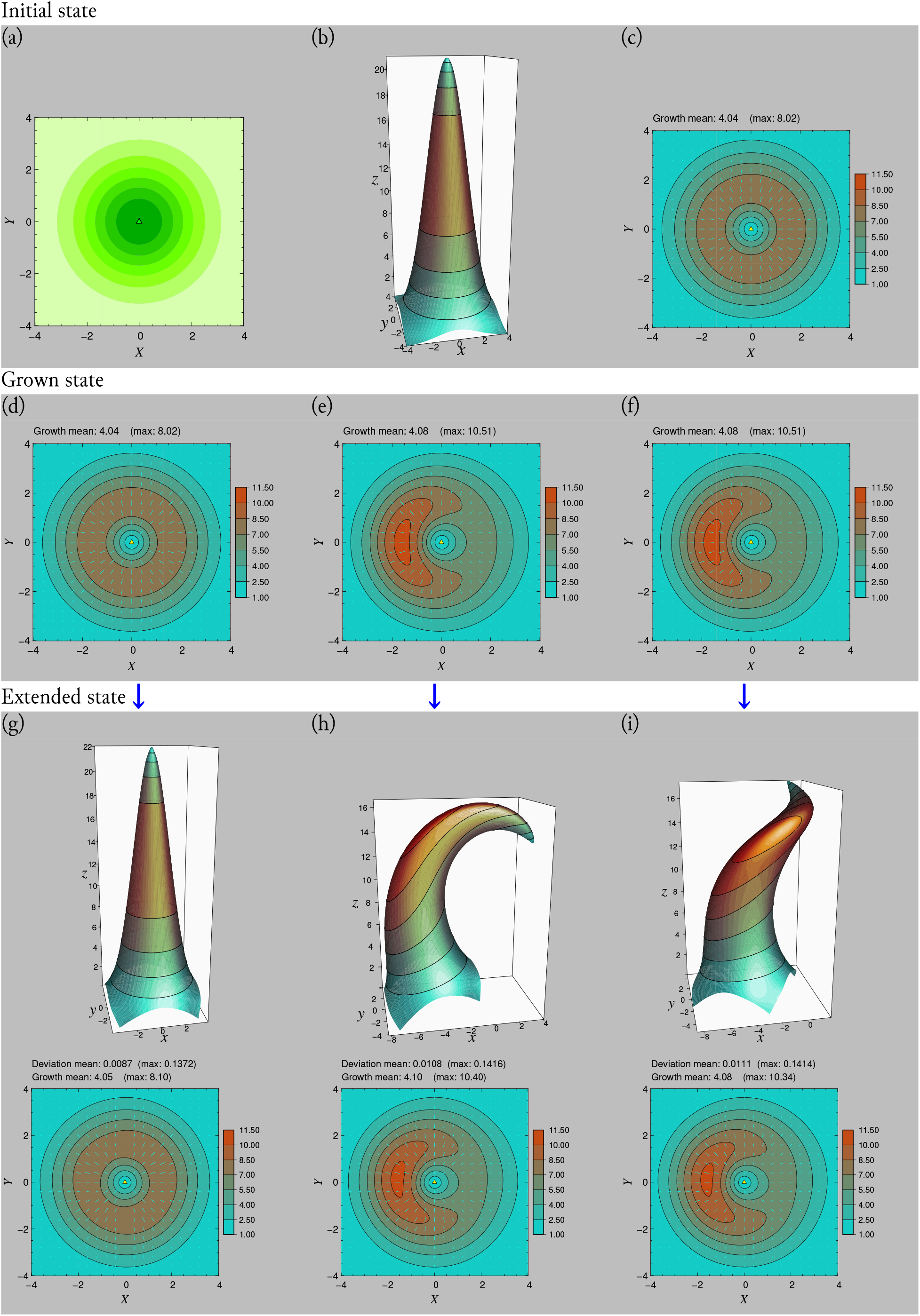
Initial state, grown state, and extended state for generation of simple horn-like shape and its modification. (a) Morphogen intensity distribution *β*(**u**) defined as an isotropic Normal distribution. (b) Basic shape for *β*(**u**), i.e., representation of morphogen intensities as heights. (c) Basic growth metric **M**_*β*_ (**u**), i.e., the growth metric calculated from the basic shape (see Appendix B). (d) Grown sheet produced from *β*(**u**) under the MGS growth model, represented with the growth metric **M**(**u**) given by equation (6). (e) Grown sheet for curving the basic shape. (f) Grown sheet for curving and twisting the basic shape. (g), (h), and (i) show extended sheets from (d), (e), and (f), respectively, in the spatial coordinates (upper panels) and the material coordinates (lower panels, represented with the extension metric **M**_ext_(**u**)). See Appendix D for how to curve and twist the basic shape.

In the present study, we investigated how complex 3D structures emulating beetle horns or mandibles can be generated through the non-uniform growth of epidermal sheets regulated by morphogen distributions. To facilitate the analysis, we adopted the MGS growth model [18], because, as described above, this model provides simple correspondence of a morphogen distribution to the shape of an extended sheet. For efficient analysis on the relationship between the growth regulation and the extended sheet shape, our analysis did not incorporate the epidermal folding and remodelling process during beetle pupation [19]. Our results show that heterochronic expression of several morphogens on the epidermal sheet can fuse different simple shapes to generate various complex shapes.

The remainder of this paper is organised as follows. Section 2 explains how the nonuniform sheet growth in the MGS growth model is described geometrically, and how the 3D shape of the extended sheet is analysed. Section 3a shows how multiple simple shapes can be fused in the extended sheet and how the overlap of expression periods among morphogens affects the shape of the extended sheet. Section 3b shows that the fusion of simple shapes can flexibly generate various 3D shapes. Finally, section 4 discusses our results in relation to previous studies.

## 2. Methods

### (a) Description of sheet growth

In the MGS growth model, the epidermal sheet grows only in the direction tangential to the sheet surface (corresponding to the surface area increase) at each position; growth in the normal direction (corresponding to the thickness increase) is not considered. Therefore, for notational simplicity, the sheet is assumed to be infinitesimally thin (although the sheet thickness was explicitly considered in the finite element method analysis for the shape of the extended sheet).

Initially, the sheet is assumed to lie on the *x*-*y* plane (*z* = 0) in the 3D spatial coordinate system **r** = (*x,y,z*)^T^, where the superscript T denotes the transposition of a matrix or vector (figure 1a). On this sheet, a two-dimensional coordinate system, **u** = (*X,Y*)^T^, is set, providing the material coordinates of the sheet, called the Lagrangian coordinate system in continuum mechanics (figure 1b). The material coordinate system stretches or contracts in the same manner as the sheet stretches or contracts. Hence, the sheet maintains the same shape in the material coordinate system with its initial state, regardless of how it has deformed in the spatial coordinate system (figures 1b, 1d, and 1f). Accordingly, the position of each microelement in the sheet is invariant in the material coordinate system.

For a microelement of the sheet at an arbitrary position **u** in the material coordinate system, its spatial coordinates at time *t* are denoted by 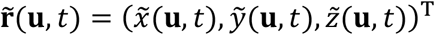. In the initial state, the (*x, y*) components of the spatial coordinates coincide with the material coordinates, satisfying the conditions 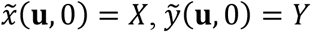, and 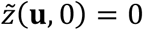.

For describing the non-uniform growth of the sheet, a short line segment from **u** = (*X,Y*)^T^ to **u** + **δu** with **δu** = (*δX, δY*)^T^ is considered in the material coordinates. Then, its ‘unstrained length’ at time *t*, denoted by *l*(**u, u** + **δu**, *t*), is defined as its length in the spatial coordinates, provided that the microelement including the line segment undergoes no stress. On this basis, the ratio of the unstrained length at time *t* to its initial length |**δu**| describes the growth amount of the sheet at **u**. Specifically, the growth amount *L*(**u, e**, *t*) at **u** along direction **e** = **δu** / |**δu**| at time *t* is given by

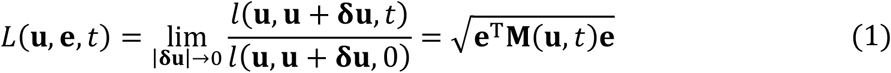

(see Appendix A.1 for the transformation), where **M**(**u**, *t*) is a two-by-two symmetric matrix given by

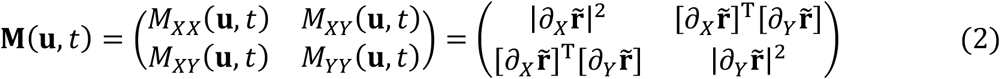

with 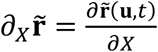 and 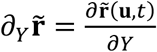. Here, **M**(**u**, *t*) is referred to as the growth metric, containing all the information on the amount of growth in all directions. For example, **e** = (1,0)^T^ gives 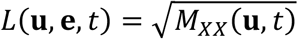, whereas **e** = (0,1)^T^ gives 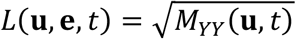. In the initial state, the growth amount is assumed to be equal to 1 in any direction, satisfying *L*(**u, e**, 0) = 1 for any **e**, meaning that **M**(**u**, 0) = **I** holds with **I** describing the identity matrix.

### (b) Sheet growth in MGS growth model

Mathematical descriptions for non-uniform sheet growth often result in formulas that make analysis difficult. By contrast, as shown below, the MGS growth model describes the non-uniform sheet growth in a considerably simple form, allowing us to obtain deep insights.

In the MGS growth model, the sheet at each position in the spatial coordinates grows in parallel with the intensity gradient of a morphogen along the sheet at that position, and its growth rate is proportional to the square of the gradient strength. The morphogen intensity at each position **u** at time *t*, denoted by *β*(**u**, *t*), is assumed to be unaffected by the sheet growth. Sheet growth is assumed to be unstrained in the sense that the actual growth of each microelement of the sheet in spatial coordinates follows the growth amount given by equation (1). Note that the morphogen intensity gradient along the sheet in the spatial coordinates can still be weakened by the sheet growth. Under these assumptions, the time change of the growth metric is independent of the sheet shape in spatial coordinates and derived as

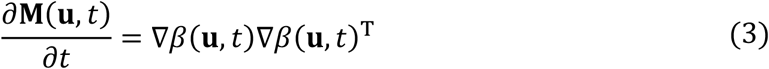

(see Appendix A for the derivation), where 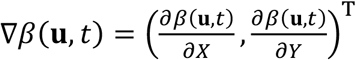 describes the gradient of *β*(**u**, *t*) in the material coordinates evaluated at **u**, and where the ∇*β*(**u**, *t*)∇*β*(**u**, *t*)^T^ is a two-by-two symmetric matrix given by

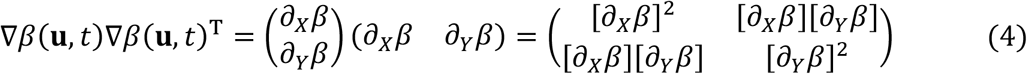

with 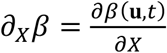 and 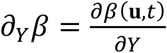. Because **e** in equation (1) does not depend on time, we see from equations (1) and (3) that

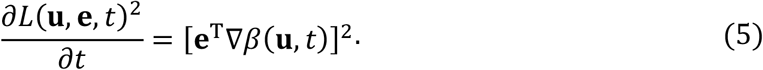

Hence, **e** being parallel to ∇*β*(**u**, *t*) gives ∂*L*(**u, e**, *t*)^2^/∂*t* = |∇*β*(**u**, *t*)|^2^ (i.e., morphogen-gradient-squared growth), whereas **e** being orthogonal to ∇*β*(**u**, *t*) gives ∂*L*(**u, e**, *t*)/∂*t* = 0.

When the morphogen distribution can be expressed as *β*(**u**, *t*) = *ρ*(*t*)*β*(**u**) with *ρ*(*t*) describing the temporal expression pattern (satisfying *ρ*(*t*) = 0 for *t* ≥ τ) for a static morphogen distribution *β*(**u**), the time integration of equation (3) gives the growth metric at time *t* = τ:

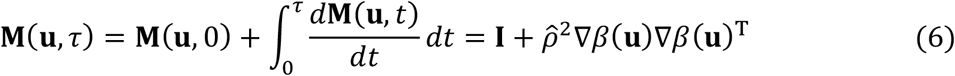

with 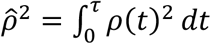. As derived in Appendix B.1, this growth metric, **M**(**u**, τ), is identical to the growth metric **M** _*β*_ (**u**, τ) attained when the sheet shape is given by

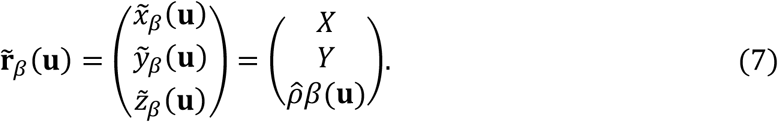

In other words, for each microelement of the sheet at its initial state 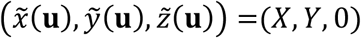, only the *z*-coordinate is raised from 0 to 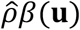 (Appendix B.1). Therefore, the grown sheet with its growth metric **M**(**u**, τ) can be extended into a smooth shape 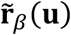 that makes the sheet unstrained (provided that the sheet is infinitesimally thin), where 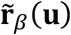 is called an isometric embedding for metric **M**(**u**, τ) in differential geometry. Hence, the MGS growth model can generate a 3D structure (from a flat epithelial sheet) corresponding to the shape of the morphogen distribution.

For convenience, *β*(**u**) is rescaled so that 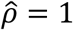 holds without loss of generality. We then refer to the 3D shape 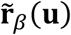 and its corresponding growth metric **M**_*β*_(**u**, τ) as the ‘basic shape’ and ‘basic growth metric’, respectively, induced from morphogen distribution *β*(**u**).

### (c) Analysis of the 3D shape of the extended sheet

To investigate the extended shape of the grown sheet, we numerically analysed the sheet deformation in spatial coordinates by means of the finite element method with the neo-Hookean elasticity assumption (using the modular finite element method [MFEM] library in C++ language [20]; see Appendix C.1 for details). The initial state of the sheet is a thin square object placed on the *x*-*y* plane (side length = 8.0, along the *x*- and *y*-axes, and thickness = 0.2 along the *z*-axis). Differently from the sheet growth process assuming unstrained growth, the extended sheet may have some strain so that the ‘extension metric’ (figure 1f), calculated from the shape of the extended sheet (Appendix B.1), may deviate from the growth metric (figure 1d) given by equation (6).

In actual moulting in arthropods, the epidermal sheet detached from the old cuticle grows and produces a new cuticle with weak bending elasticity, forming wrinkled primordia [16]. After removing the old cuticle, these primordia are then unfolded by water pressure into a smooth shape with stronger elasticity [21]. However, such a wrinkled state causes difficulty in calculation of the sheet deformation. Therefore, to avoid the wrinkled state in our analysis, we assumed that the sheet always had sufficiently strong elasticity to allow its steady extension by water pressure into a smooth 3D shape.

When the morphogen distribution *β*(**u**) is defined as a normal distribution (figure 2a), the grown sheet (figure 2d) is extended into a 3D shape (figure 2g) similar to its corresponding basic shape (figure 2b). The extended sheet can be curved (figure 2h) or twisted (figure 2i) by assuming additional morphogens that modify the degree and direction of local sheet growth (see Appendix D for details).

We also analysed the shapes of the extended sheets that mimic the head and thorax of beetles with horns (see Appendix C.2 for details). Although the finite element method can calculate sheet deformation under realistic physical assumptions, the calculation may not be stable when the sheet is considerably thin as in the case of insect exoskeletons. Hence, in this analysis, we used the one-layer vertex-spring method, which is expected to be more stable for thin sheets as it does not require an explicit description of the sheet thickness.

## 3. Results

In this section, we first show that the MGS growth model can flexibly fuse multiple basic shapes through the heterochronic expression of multiple morphogen distributions on the sheet. We then show that various 3D shapes of extended sheets can be generated by fusing basic shapes.

### (a) Fusion of basic shapes

Multiple basic shapes can be fused by expression of multiple morphogen distributions on the same sheet. Specifically, we can fuse the basic shapes of two static morphogen distributions, denoted by *β*_1_(**u**) and *β*_2_(**u**), by defining *β*(**u**, *t*) as

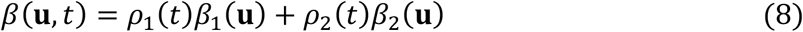

with *ρ*_1_(*t*) and *ρ*_2_(*t*) describing the temporal expression patterns for *β*_1_(**u**) and *β*_2_(**u**), respectively, and where both expressions end by time *t* = τ (i.e., *ρ*_1_(*t*) = 0 and *ρ*_2_(*t*) = 0 for *t* ≥ τ) and satisfy 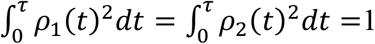. The final growth metric at *t* = τ is then given by

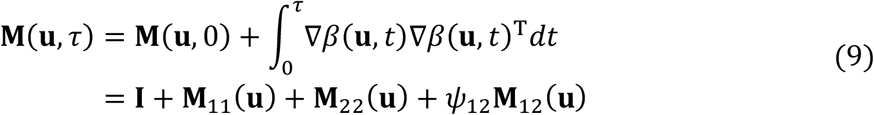

with

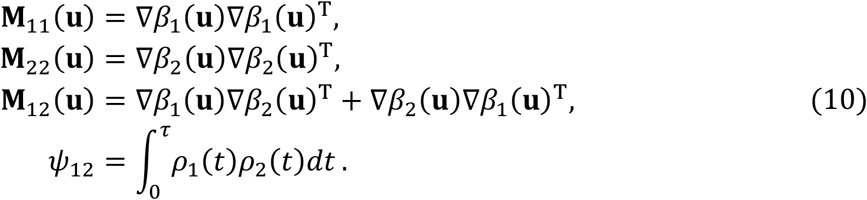

Hence, if we modify *ρ*_1_(*t*) and *ρ*_2_(*t*), keeping their temporal overlap *ψ*_12_ unchanged, then **M**(**u**, τ) is also unchanged. In particular, when *ψ*_12_ = 0, switching of the expression order between *β*_1_(**u**) and *β*_2_(**u**) does not affect **M**(**u**, τ).

Figure 3 shows how *ψ*_12_ affects the shape of the extended sheet when *β*_1_(**u**) and *β*_2_(**u**) are given by wide (green gradient in panel (a)) and narrow normal distributions (blue contours in panel (a)), respectively. If *ρ*_1_(*t*) = *ρ*_2_(*t*), then *ψ*_12_ attains its maximum value of 1. In this case, the growth metric (figure 3e) induced from Eq. (8) is identical to that induced from a single morphogen distribution defined by *β*(**u**) = *β*_1_(**u**) + *β*_)_(**u**) (figure 3c). Therefore, the shape of the extended sheet (figure 3h) resembles the basic shape for *β*(**u**) = *β*_1_(**u**) + *β*_2_(**u**) (figure 3d).

**Figure 3.**
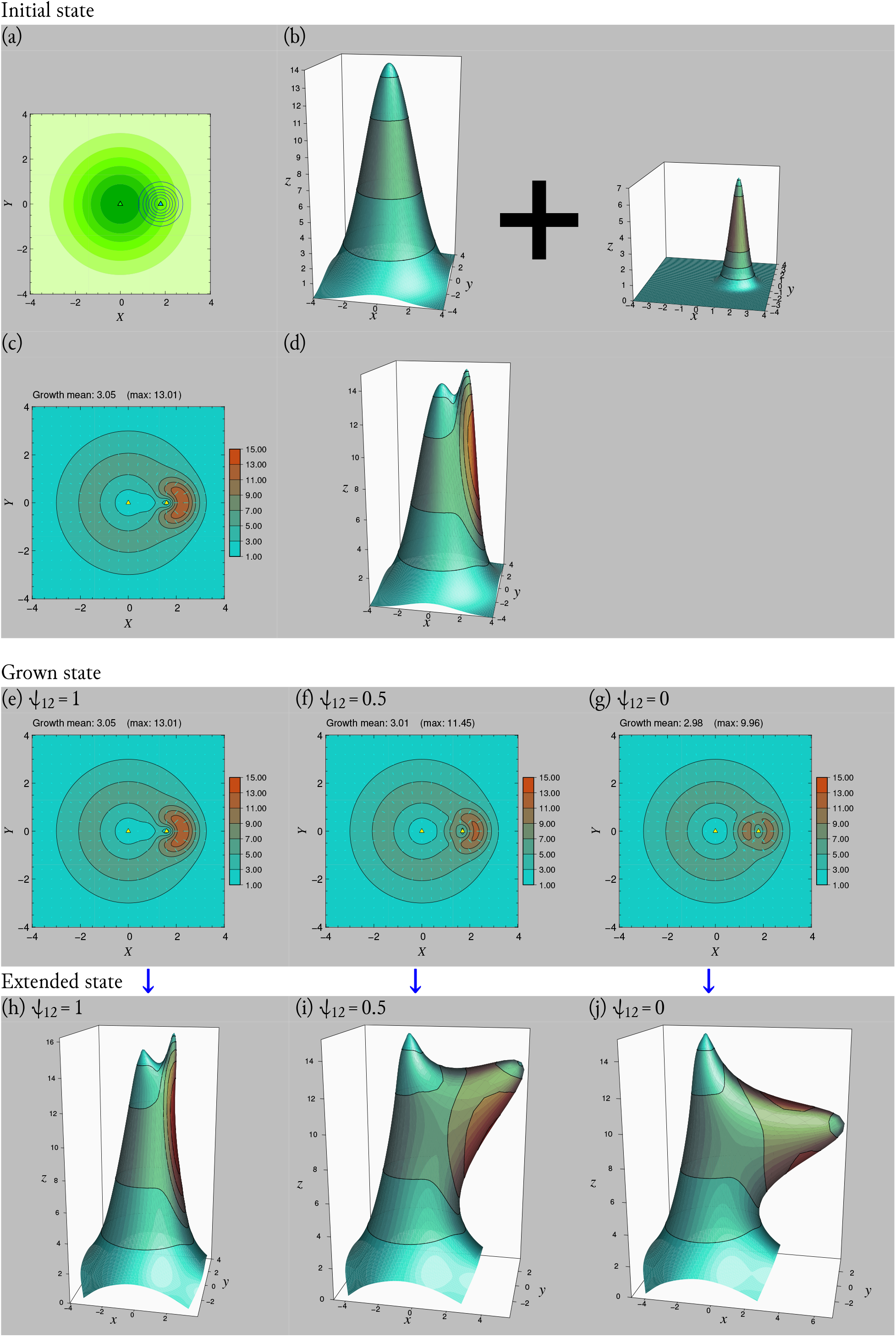
Fusion of two basic shapes by heterochronic expression of two morphogen distributions. (a) Two morphogen distributions *β*_1_(**u**) and *β*_2_(**u**) plotted with green gradient and blue contours, respectively, with small triangles indicating their maximums. (b) Basic shapes for *β*_1_(**u**) (left) and *β*_2_(**u**) (right). (c) Basic growth metric for *β*_1_(**u**) + *β*_2_(**u**). (d) Basic shape for *β*_1_(**u**) + *β*_2_(**u**). (e), (f), and (g) show grown sheets for *ψ*_12_ = 1, *ψ*_12_ = 0.5, and *ψ*_12_ = 0, respectively. (h), (i), and (j) show extended sheets for *ψ*_12_ = 1, *ψ*_12_ = 0.5, and *ψ*_12_ = 0, respectively. Plotting format: panel (a) is the same as those shown in figure 2a; panels (b), (d), and (h-j) are the same as those shown in figure 2g-i; panels (e-g) are the same as those shown in figure 2d-f. See Appendix C.1 for morphogen distributions and parameters.

By contrast, if *ρ*_1_(*t*) and *ρ*_2_(*t*) have no temporal overlap, *ψ*_12_ attains its minimum value of 0. In this case, orientations of the substructures induced from *β*_1_(**u**) (left object in figure 3b) and *β*_2_(**u**) (right object in figure 3b) tend to be orthogonal to each other in the extended sheet (figure 3j). Note that the sheet growth induced from *β*_1_(**u**) at the position of *β*_2_(**u**) broadens the shape of the substructure induced from *β*_2_(**u**) (figure 3j) in comparison with its basic shape (right object in figure 3b), in accord with the mathematical derivation in Appendix B.2. When *ψ*_12_ has an intermediate value between 0 and 1, the orientations of the substructures tend to have an angle between parallel and orthogonal (figure 3i).

Analogously, we can fuse the basic shapes for an arbitrary number of morphogen distributions. For example, figure 4 shows the fusion of three basic shapes specified by three morphogen distributions: *β*_1_(**u**) (green gradations in figures 4a–4c), *β*_2_(**u**) (blue contour), and *β*_3_(**u**) (red contour), where only *ψ*_23_ is non-zero and affects the position of *β*_3_(**u**) as well as additional curving and twisting (see Appendix D for the derivation of a relationship analogous to equation (9) under additional curving and twisting). The resulting shape variation depending on *ψ*_23_ may resemble the trimorphism of male mandibles of stag beetle species in the genus *Dorcus*, depending on their body sizes [22,23].

**Figure 4.**
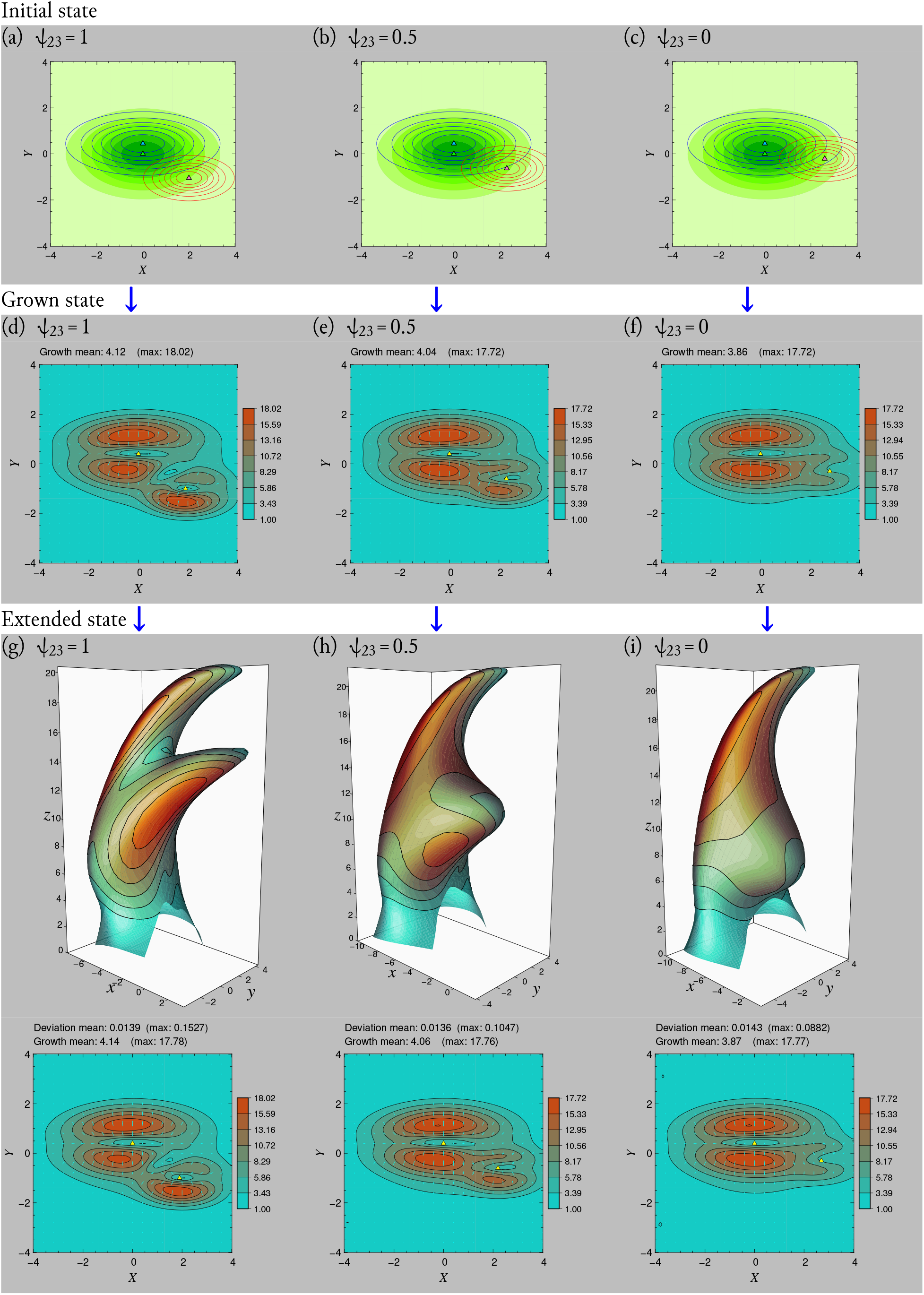
Extended sheets mimicking three mandible types of stag beetles depending on their body sizes (large-, middle-, and small-types). The extended sheets are generated by fusing the basic shapes of three morphogen distributions, *β*_1_(**u**), *β*_2_(**u**, *ψ*_23_), and *β*_3_(**u**, *ψ*_23_), where only *β*_2_ and *β*_3_ have temporal overlap in their expression (i.e., *ψ*_12_ = *ψ*_13_ = 0 and 0 ≤ *ψ*_23_ ≤ 1). (a), (b), and (c) show morphogen distributions for the large-type (*ψ*_23_ = 1), middle-type (*ψ*_23_ = 0.5), and small-type (*ψ*_23_ = 0). Morphogen distributions *β*_1_, *β*_2_, and *β*_3_ are plotted respectively as green gradient, blue contours, and red contours, with small triangles indicating their maximums. (d), (e), and (f) show grown sheets for the large-type (*ψ*_23_ = 1), middle-type (*ψ*_23_ = 0.5), and small-type (*ψ*_23_ = 0). (g), (h), and (i) show extended sheets (upper panels) and the corresponding extension metrics (lower panels) for the large-type (*ψ*_23_ = 1), middle-type (*ψ*_23_ = 0.5), and small-type (*ψ*_23_ = 0). Plotting formats are the same as those shown in figure 3. See Appendix C.1 for morphogen distributions and parameters.

### (b) Morphogenesis of complex shapes

Fusion of multiple basic shapes with additional curving and twisting can generate various complex 3D shapes for the extended sheet (figure 5). Note that these complex shapes are generated from comparably much simpler settings for morphogen distributions given by normal distributions or their linear combinations (Appendix C.1), and which may provide an efficient genetic coding for various 3D morphologies as discussed in the Discussion (b).

**Figure 5.**
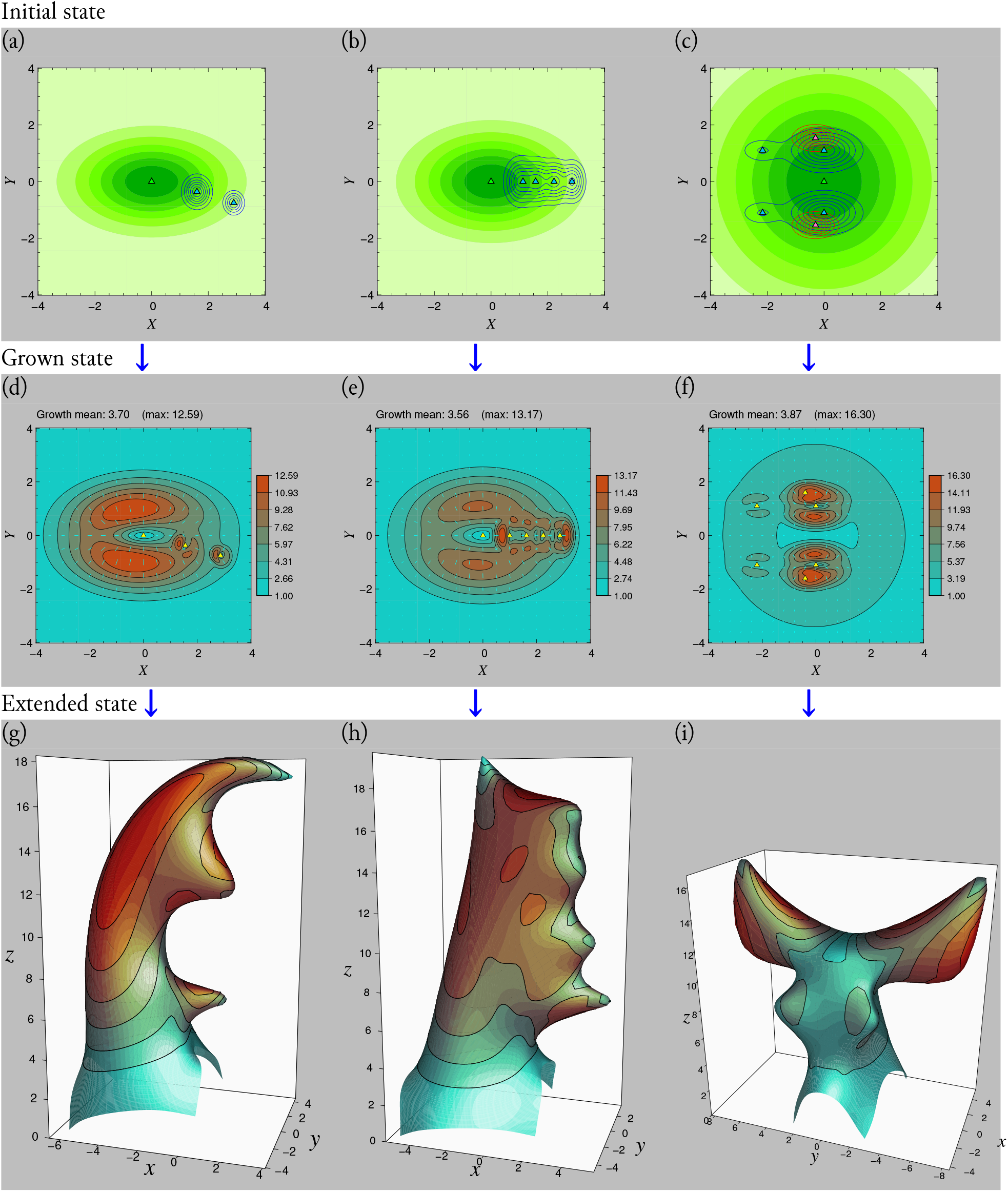
Extended sheets of complex 3D shapes generated by fusing multiple basic shapes. (a-c) Morphogen distributions. (d-f) Grown sheets represented with growth metrics. (g-i) Extended sheets. Plotting formats are the same as those shown in figures 3 and 4. See Appendix C.1 for morphogen distributions and parameters.

Figures 6–8 show the extended sheets obtained with the vertex-spring method, where the morphogen distributions and their combinations are designed for mimicking heads and thoraxes of beetles with horns: figure 6 mimics genus *Dynastes* (e.g., *D. hercules*, especially *Goku-buto* (ultra thick in Japanese) breeding strain [24]) and genus *Xylotrupes* (e.g., *X. pubescens*), figure 7 mimics genus *Chalcosoma* (e.g., *C. chiron*), and figure 8 mimics genus *Sulcophanaeus* (e.g., *S. faunus*). Analogously to figure 5, these 3D shapes were generated by fusing multiple basic shapes with additional curving and twisting, where all assumed morphogen distributions are normal distributions or their linear combinations.

**Figure 6.**
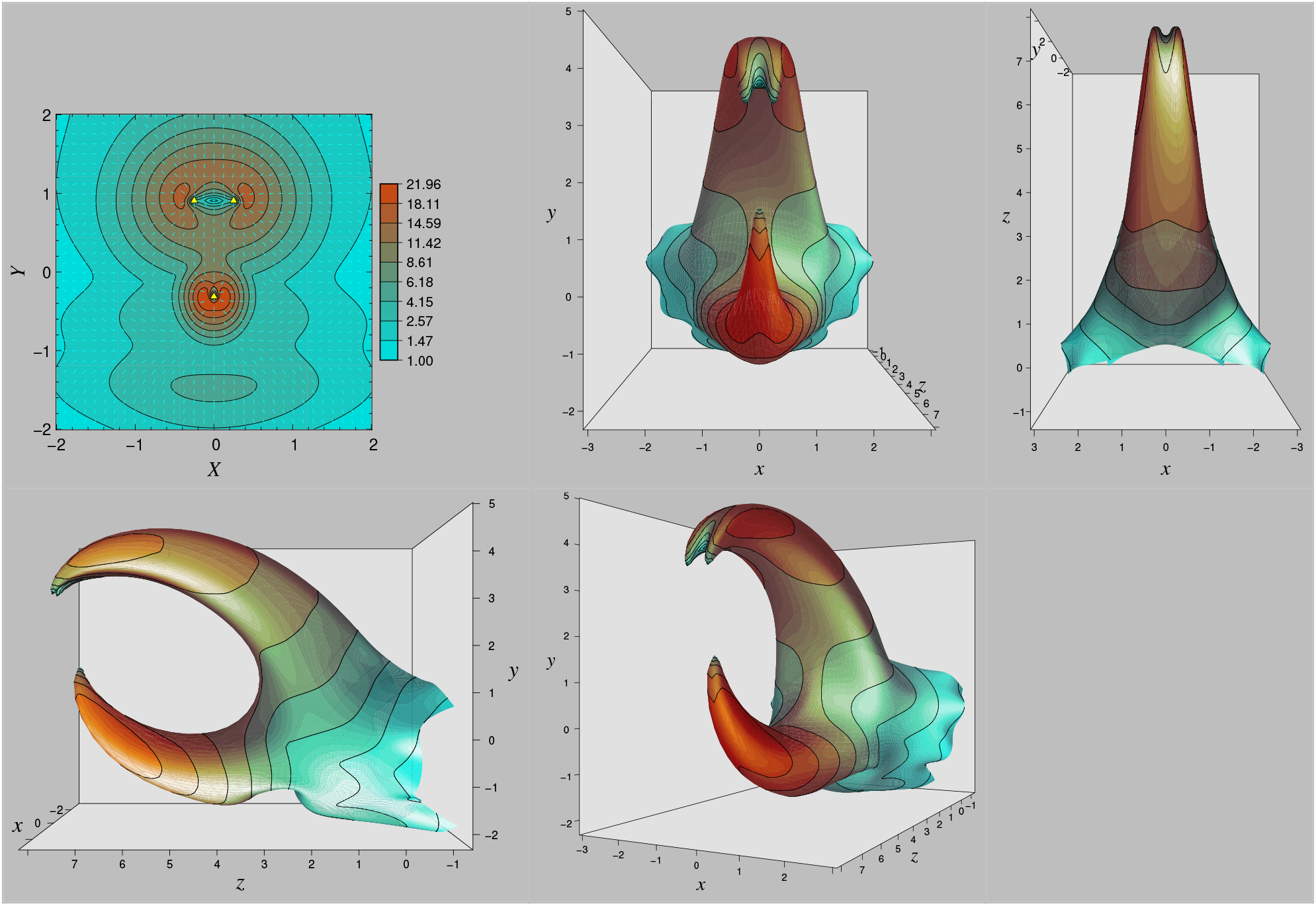
Extended sheet mimicking the head and thorax of beetles in the genus *Dynastes* (e.g., *D. hercules*, especially *Goku-buto* (ultra thick in Japanese) breeding strains [24]) and genus *Xylotrupes* (e.g., *X. florensis*). The upper left panel shows the grown sheet represented with the growth metric, and the other panels show its extended shape viewed from different perspectives. See Appendix C.2 for morphogen distributions and parameters.

**Figure 7.**
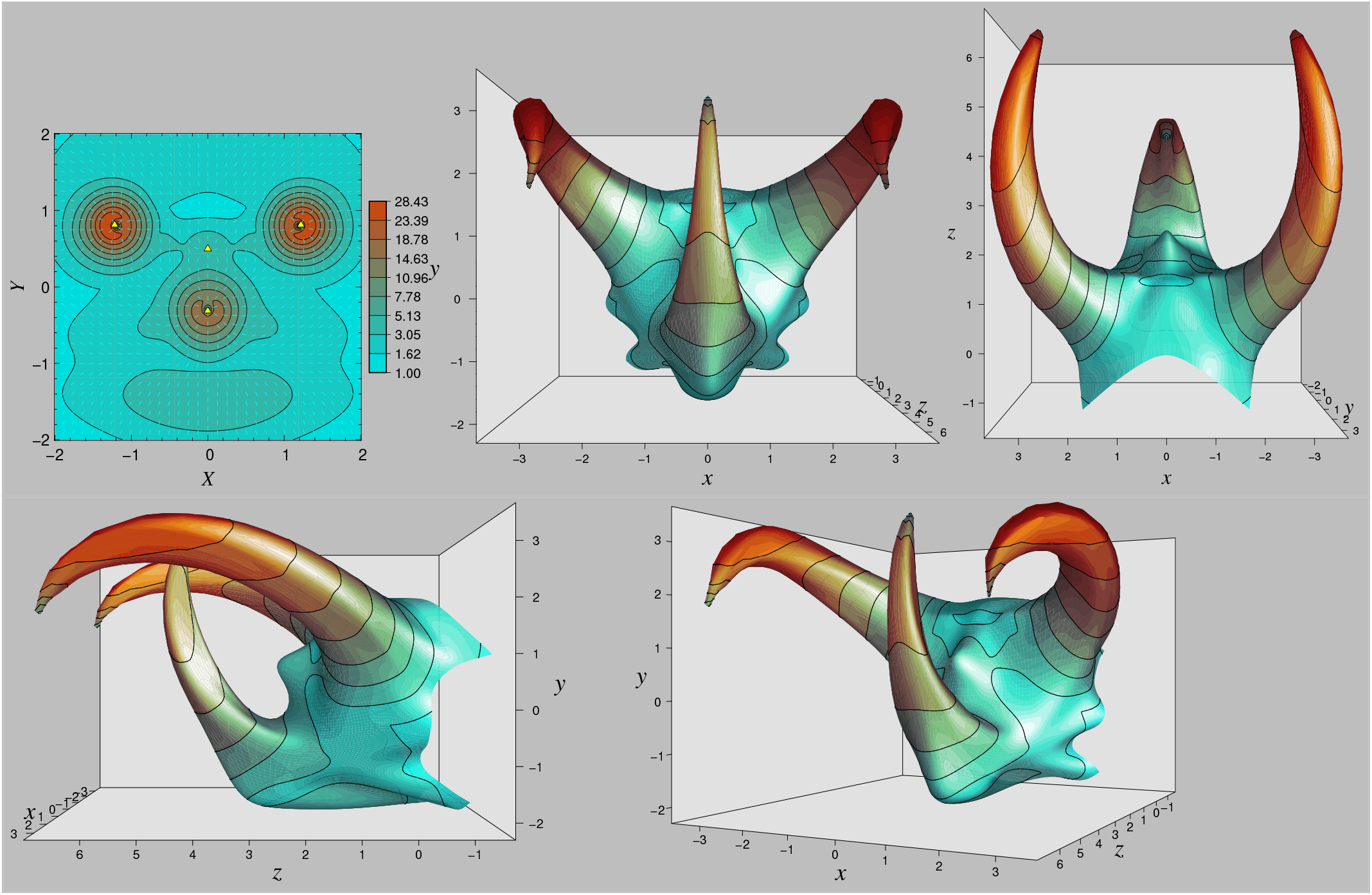
Extended sheet mimicking the head and thorax of beetles in the genus *Chalcosoma* (e.g., *C. caucasus*). Plotting formats are the same as in figure 6. See Appendix C.2 for morphogen distributions and parameters.

**Figure 8.**
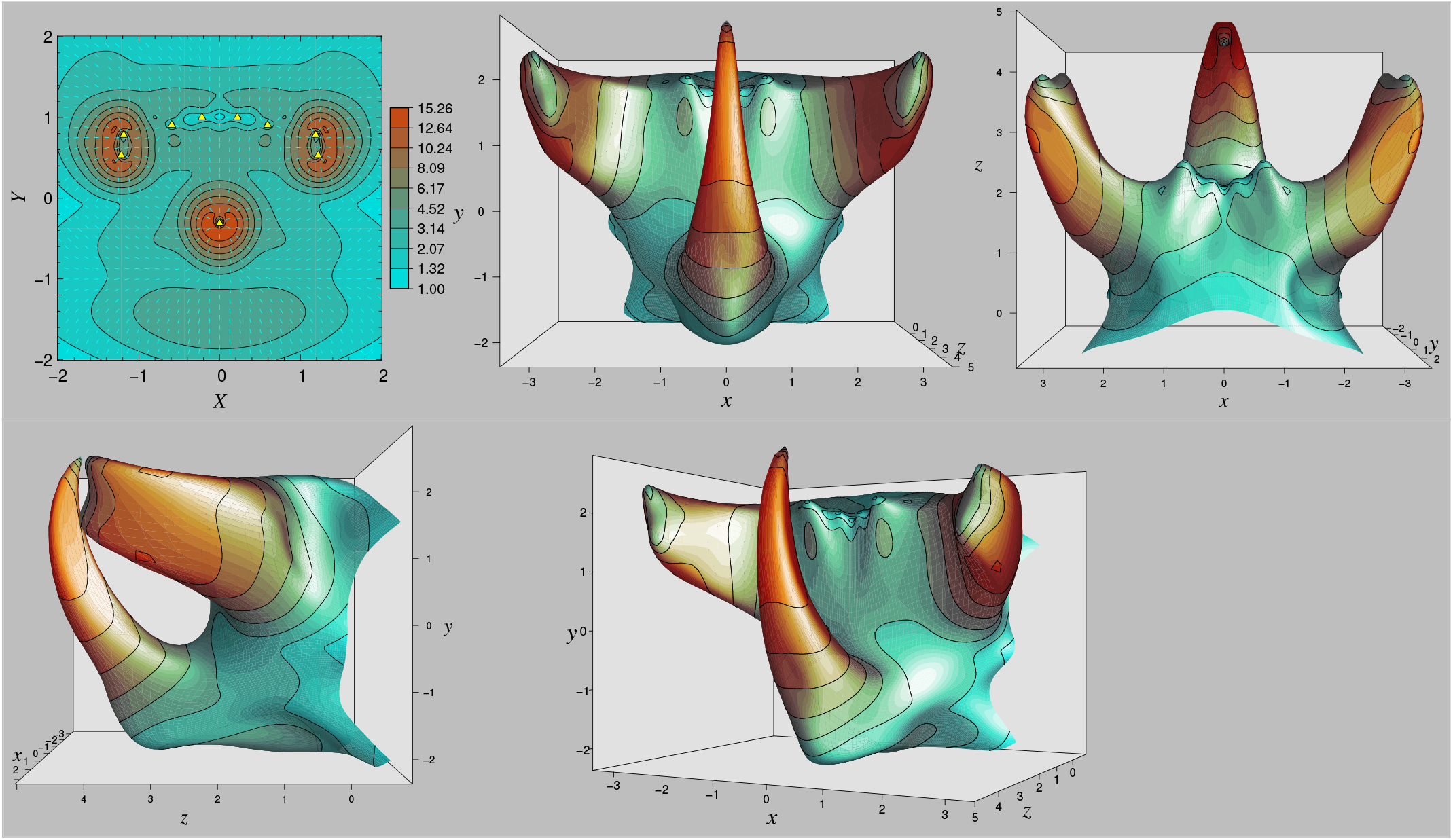
Extended sheet mimicking the head and thorax of beetles in the genus *Sulcophanaeus* (e.g., *S. faunus*). Plotting formats are the same as in figure 6. See Appendix C.2 for morphogen distributions and parameters.

## 4. Discussion

### (a) Growth regulation of epithelial sheet and developmental constraints

The present study showed that the non-uniform growth in the MGS growth model [18] can transform the planar epidermal sheet into various complex shapes emulating the horn shapes of actual beetles. The MGS growth model is a deductively devised minimal model for achieving smooth 3D shapes, rather than representing the actual process of arthropod morphogenesis. Nevertheless, the generated structures not only resemble the actual horns of beetles as a whole but also share some local shapes. For example, in the morphogenesis of these horn-like shapes, the maximum sheet growth (corresponding to the maximum morphogen gradient) is attained in the middle part of the horn and oriented toward the apex (figure 2d), following the empirically observed sheet growth for horn morphogenesis in beetles [9]. Moreover, in the morphogenesis of complex shapes, the generated structures deviate spontaneously from the shapes of morphogen distributions, and the deviations bring local distortions that are similar to that of actual horns. In particular, the curving and twisting of the horns shown in figures 6–8 spontaneously transforms their cross-sections from round shapes into angular ones that resemble actual beetle horns [4].

As the angular cross-sections, spontaneous deformation that deviates from the basic shape of a morphogen distribution is caused by not being able to unfold a sheet as much as it grew. The extended shape of a grown sheet corresponds to the minimisation of the sum of stretching energy (tangential to the sheet surface) and bending energy (orthogonal to the surface) over the sheet, under a negligibly low water pressure. Provided that the sheet is sufficiently thin, the stretching energy has the dominant contribution to this minimisation. If the extended sheet has a shape such that the local sheet growth measured in spatial coordinates exactly follows the growth metric derived from morphogen distributions (equation (6)), then such a shape is called an isometric embedding for the growth metric, giving zero stretching energy over the sheet.

For a sheet grown in the MGS growth model under a single morphogen distribution without additional curves and twists, the basic shape of the morphogen distribution gives the isometric embedding of the sheet, and hence the extended sheet can have a similar shape to the basic shape. By contrast, for more complex shapes generated from multiple morphogens with added curves and twists, neither basic shapes of the morphogen distributions nor their linear combinations give isometric embeddings, and hence, the shape of the extended sheet can have significant local deviations from the basic shapes. Such deviations cause further deformation of the extended sheet as a byproduct of the global minimisation of local elastic energies. The angular cross-section of the central horn in figure 7 and the concavity near the base of the left and right horns in figure 8 can be interpreted as examples of such byproduct shapes. The byproduct shapes via deviating from the shape of morphogen distributions might be regarded as developmental constraints that induce common morphological features shared by arthropods, and our model might represent some essential elements for such constraints.

Since the byproduct shapes are formed through the minimisation of elastic energies, they will contribute to the structure-mechanical stability of appendages. In particular, the horns of beetles as a weapon in male-male competition require high stability, and their angular cross-sections would be an adaptation for stability [4]. Notably, a rhinoceros beetle generates the angular shape of its horn by remodelling (i.e., adhesion and shrinkage) of the horn with a round cross section developed through pupation [19]. Our results imply that the horn before the remodelling might already be oriented to have an angular shape.

### (b) Modularity and evolution of exoskeletal morphogenesis

Even in a complex shape made by fusing multiple basic shapes under the MGS growth model, the substructure corresponding to each basic shape tends to keep its basic shape without disturbed by fusing other shapes. Notably, the size and shape of each basic shape can be changed separately by modifying the corresponding morphogen distribution. Moreover, the relative orientations among the substructures can also be changed by modifying the temporal overlaps among morphogen expressions. These properties of the MGS growth model imply that substructures of appendages, such as multiple horns, spines, or branches of horns, have developmental independency bringing modularity to exoskeletal appendages. This modularity explains why our simple growth model can easily generate complex structures.

Modularity results in developmental stability and evolutionary robustness against mutations. Even if an exoskeletal structure is structure-mechanically stable, mutations in the epidermal sheet growth may collapse the structure, because it is not warranted that an arbitrary grown sheet disturbed by mutations can be extended into a smooth 3D shape with structure-mechanical stability (in potential association with the increase of moulting failure observed in successive breeding of stag beetles for individuals with thicker mandibles [25]). Such collapse of exoskeletal structure caused by mutation will be effectively reduced, if an exoskeletal appendage has modularity allowing its parts to develop independently so that a mutation in one part may have little interference with other parts. In particular, under the MGS growth model, increasing/decreasing morphogen expression tends to induce only the scaling, addition, or removal of appendages (such as spines and branches) or modifications (such as curving and twisting). Moreover, changes of the expression timings of morphogens tend to modify the aspect ratios and relative orientations of the substructures without breaking down the entire structure.

Although whether the MGS growth model corresponds to actual morphogenesis remains to be examined, our study implies that even simple regulation such as the MGS growth model can afford developmental modularity. Such modularity will potentially bring developmental stability and ease of adding modifications to the morphogenesis of exoskeletal appendages, it may contribute to explaining the rapid and dramatic evolution of beetle horns with extreme shapes and diversity. The horns of beetles have been evolving rapidly and dramatically despite their extreme sizes and elaborate shapes; they maintain structure-mechanical strength and developmental plasticity as weapons for males with good physiques [2]. This ambivalent feature between evolvability and developmental stability is a classical paradox in evolutionary developmental biology. One way to resolve this paradox is the modularity of developmental units at various scales [5]. Modularity brought by a simple growth regulation such as the MGS growth model may allow the horns of beetles to evolve rapidly without losing their stability. More fundamentally, developmental systems that enhance compatibility among mutations through modular morphogenesis, such as the MGS growth model, would be favoured by selection as they bring high developmental stability. Such systems might have been acquired by arthropods in their early evolution, paving an efficient way to evolve further novel and stable exoskeletons. Once developmental systems have evolved to generate highly stable exoskeletons, they can be hardened to that extent, thereby increasing performance in locomotion, attacks on prey, and defence against enemies. Thus, the emergence and rapid diversification of arthropods with hard exoskeletons during the Cambrian Explosion [26,27] may be related to the evolutionary emergence of developmental systems that efficiently produce novel and stable exoskeletons through modular morphogenesis.

### (c) Limitations and future directions in connection with relevant studies

Because the MGS growth model is a deductively devised minimal model that relates shapes of extended sheets with hypothesised morphogen distributions, its relationship with actual molecules and biochemical processes must be examined carefully. For example, empirical studies on the morphogenesis of beetle horns have shown that the grown epithelial sheet has an intricate wrinkled structure [16,28], which unfolds and extends to form the horn through moulting to the pupa [9]. The wrinkle pattern is related to the direction and amount of local sheet growth; however, it remains unclear whether (i) wrinkle formation *induces* sheet growth or (ii) it *is induced by* sheet growth. Our present study shows that it is possible to generate extended sheets of various shapes without wrinkle formation. The results support the possibility of (ii) but do not rule out the possibility of (i). Hence, it is necessary to analyse the sheet deformation in spatial coordinates as a continuous dynamics that involves both the wrinkle structure formation through the sheet growth and its unfolding into a smooth 3D shape.

Although the MGS growth model assumes an unstrained growth of the epidermal sheet, actual sheet growth and its folding can cause mechanical stress on the sheet. This mechanical stress would be a signal for each cell to determine the appropriate growth amounts. Further investigation is needed to clarify the distribution of the mechanical stress and how the growth of each microelement responds to it (equations (6) or (9)) to generate a harmonious overall 3D shape [8]. In our study, the head and thorax of each horned beetle were mimicked by a single sheet without separation (figures 6–8). Such a global regulation of morphogenesis across multiple body segments is observed in some arthropods [29], and which might be essential for the harmonious morphogenesis of the entire body in terms of functional mechanics.

The MGS growth model is one of growth regulation models for cell sheet deformation. Since it assumes a single type of layered sheet, the growth metric corresponding to a protrusion (extending outward from the body surface) may be identical to that corresponding to a depression (sinking inward from the body surface). Hence, whether a protrusion or depression is formed depends on the initial growth direction and surrounding structures. By contrast, some growth models proposed for plant leaves and petals assume multi-layered sheets such that the 3D shape of the extended sheet can be uniquely determined under a given growth metric for each layer [30]. Even in morphogenesis of a hard exoskeleton like those of beetles and crabs, such a multi-layered sheet may work for orientating the sheet extension, as long as only one layer hardens afterwards (otherwise slight differences in the growth metric among hardened layers at the same position of the sheet can produce a large elastic energy that may cause structure-mechanical instability).

## Conclusion

We showed that the MGS growth model allows an epidermal sheet to grow into various complex structures emulating arthropods’ appendages including beetle horns, by fusion of multiple basic shapes controlled separately by different morphogen distributions. This suggests that such a developmental system can bring modularity and developmental stability to the morphogenesis of exoskeletal appendages and provide insights into the diversity of arthropod morphology.

## Acknowledgements

H.C.I. thanks Tomonari Kaji for valuable discussions on arthropod morphogenesis. H.C.I. also thanks David H. Munro for developing a tool for numerical analysis and visualisation, named Yorick, and distributing it for free; all figures in this paper were produced with Yorick.

## Appendix A: Derivation of equation (3)

Here we derive equation (3) in the main text in a more general form than [18].

### A.1 Description of non-uniform sheet growth

Since the microelement including an arbitrary position **u** is assumed to be unstrained in the spatial coordinates, the square of the unstrained length *l*(**u, u** + **δu**, *t*) for a short line segment from **u** = (*X,Y*)^T^ to **u** + **δu** = (*X* + *δX,Y* + *δY*)^T^ is expressed as

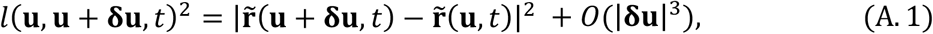

where *O*(|**δu**|^3^) contains the third- and higher-order terms of **δu**, so that there exists a constant *C* such that 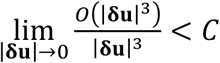 holds. To transform this equation, we expand 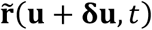 around **u** as

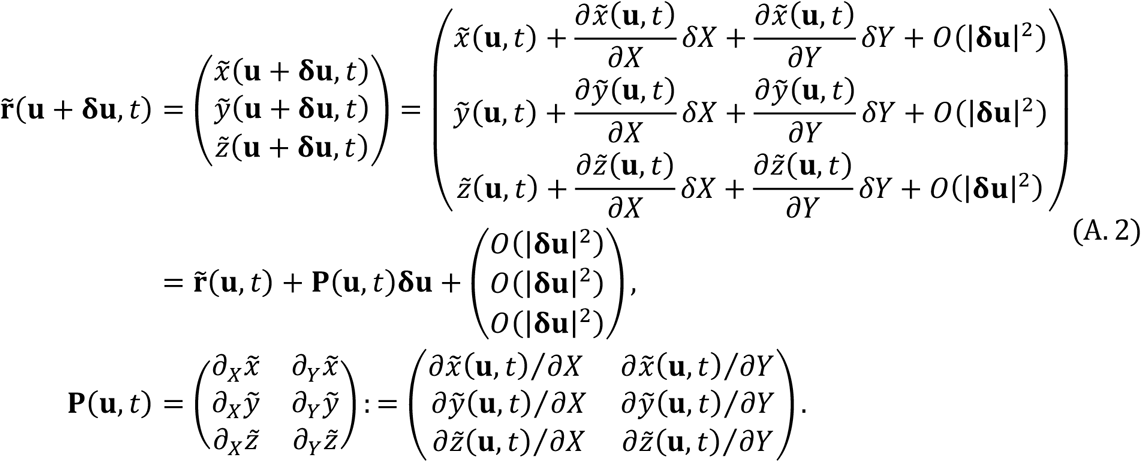

Substitution of equation (A.2) into (A.1) gives

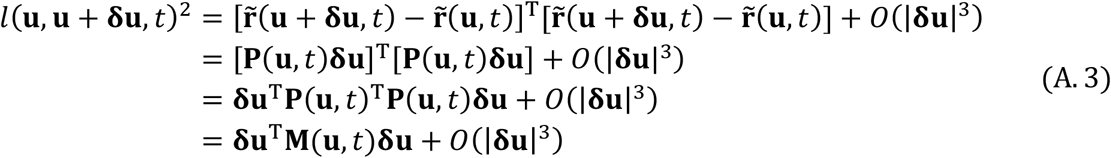

with

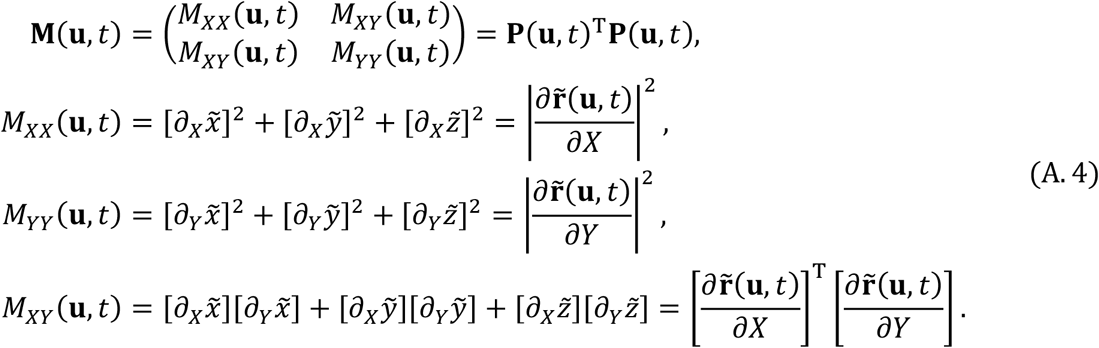

Then, the growth amount for the line segment from the initial state to time *t*, is defined by *l*(**u, u** + **δu**, *t*)/*l*(**u, u** + **δu**, 0), and which for an infinitesimally short line segment (i.e., |**δu**| → 0) describes the growth amount for the sheet at position **u** along direction **e** = **δu** / |**δu**| at time *t*:

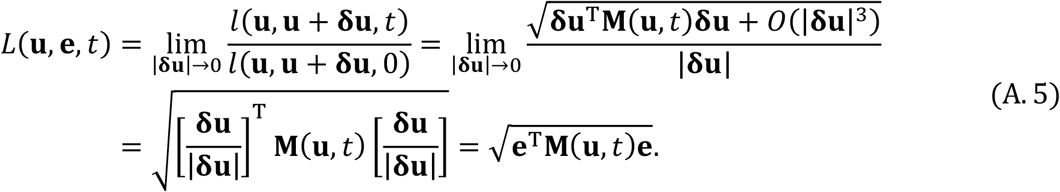

### A.2 Locally geodesic coordinate system

To describe the morphogen intensity gradient along the sheet in the spatial coordinates, we introduce a local two-dimensional coordinate system **u**^∘^ = (*X*^∘^, *Y*^∘^)^*T*^ on the sheet around each focal position 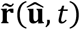 corresponding to 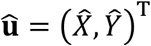. For notational simplicity, we assume **û** = **0** without loss of generality and drop the arguments “(**û**, *t*)” from variables evaluated at **û** and *t*. Since the sheet growth in the spatial coordinates is assumed to be unstrained, we can adjust coordinate system **u**^∘^ so that the unstrained length from **u** to **u** + **δu**, corresponding to the length from **u**^∘^ to **u**^∘^ + **δu**^∘^, is expressed as 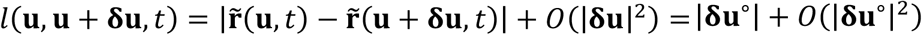 for sufficiently small |**u**| and |**δu**|. Such a coordinate system is called the locally geodesic coordinate system [31], and in which **δu** is expressed as **δu**^∘^ satisfying

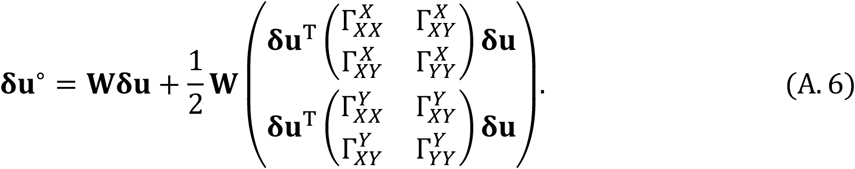

Here, **W** is a two-by-two matrix obtained by the diagonalisation of **M** as

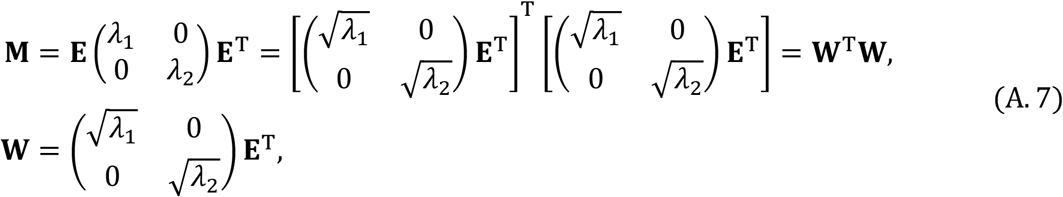

where λ_1_ and λ_2_ are the two eigenvalues of **M** with their eigenvectors **v**_1_ and **v**_2_ being combined into the matrix **E** = (**v**_1_, **v**_2_). Note that all Γs are negligible because all terms that belong to *O*(|**δu**|^2^) in equation (A.6) vanish when the limit |**δu**| → 0 is taken afterwards. (Γs are called the Christoffel symbols of the second kind in differential geometry of surfaces, and which are adjusted first derivatives of the growth metric, defined by 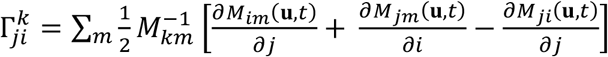 with *i, j, k, m* ∈ {*X,Y*} and 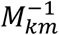 describing the (*k, m*) component of **M**^−1^ [31]). Therefore, for convenience, we transform equation (A.6) into

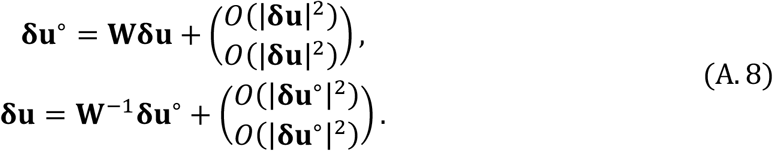

### A.3 Unidirectional and isotropic sheet growth

In the locally geodesic coordinates, **u**^∘^ = (*X*^∘^, *Y*^∘^)^*T*^, we consider both of unidirectional growth and isotropic growth for the sheet with their rates denoted by *η* and ζ, respectively. We describe the direction of the unidirectional growth at time *t* by a unit vector 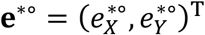. For convenience, we introduce the orthogonal unit vector to **e**^*∘^, given by **n** 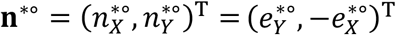. Then, we can decompose **δu**^∘^ into the element along with **e**^*∘^ and that along with **n**^*∘^:

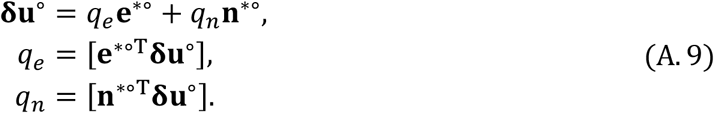

From time *t* to time *t*^′^ = *t* + Δ*t*, the *q*_D_ and *q*_E_ become 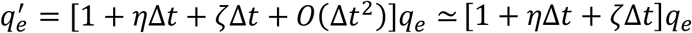 and 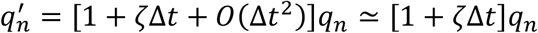, and which approximately gives

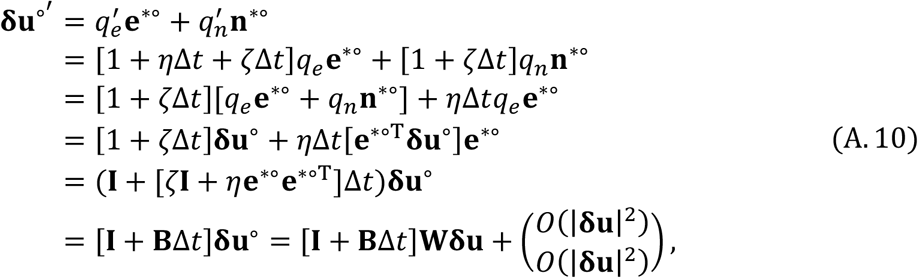

where **B** = ζ**I** + *η***e**^*∘^**e**^*∘*T*^. Hence, the growth amount at **û** = **0** along with **e** = **δu** / |**δu**| at time *t*^′^ is transformed as

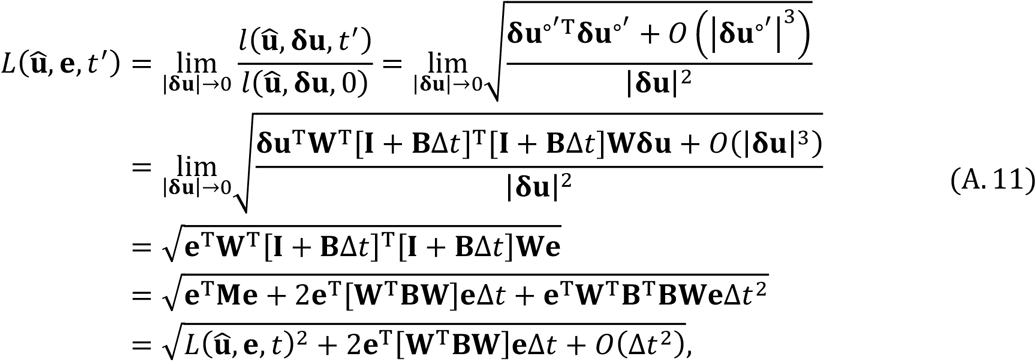

where **B**^T^ = [ζ**I** + *η***e**^*∘^**e**^*∘T^]^T^ = ζ**I** + *η***e**^*∘^**e**^*∘T^ = **B** is used. From equations (A.11) and (A.7) we get

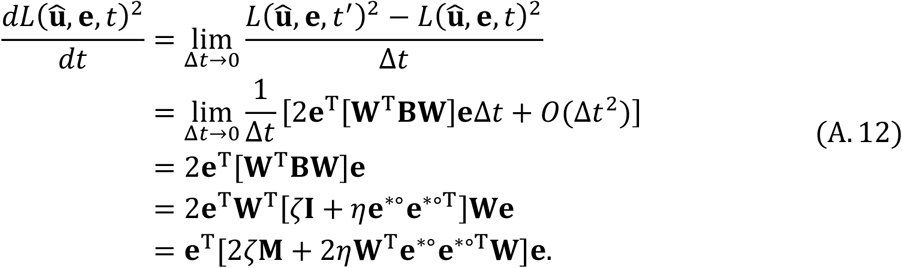

Since **e** is invariant against time, the following relationship holds good:

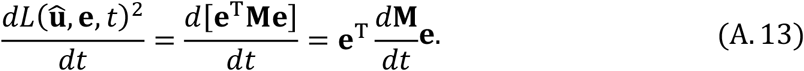

Because equations (A.12) and (A.13) both hold for an arbitrary **e**, we obtain

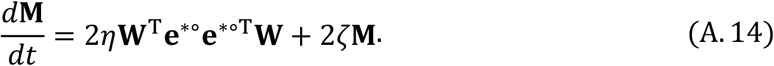

#### A.4. Sheet growth regulation by morphogen intensity gradient

Now we assume that the direction of the unidirectional growth, **e**^*∘^, is given by the gradient ∇^∘^*β* of morphogen intensity *β*(**u**) evaluated at **û** = **0** in the locally geodesic coordinates, **u**^∘^:

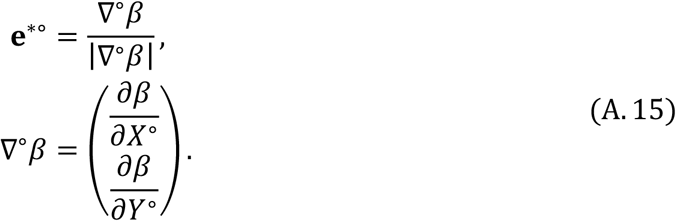

Since equation (A.8) gives **δu**^T^ = **δu**^∘*T*^ **W**^−1T^ for infinitesimally small |**δu**^∘^|, we see that **W**^−1T^ can be expressed as

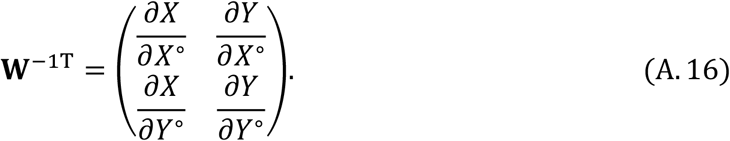

By using equation (A.16), we transform ∇^∘^*β* as

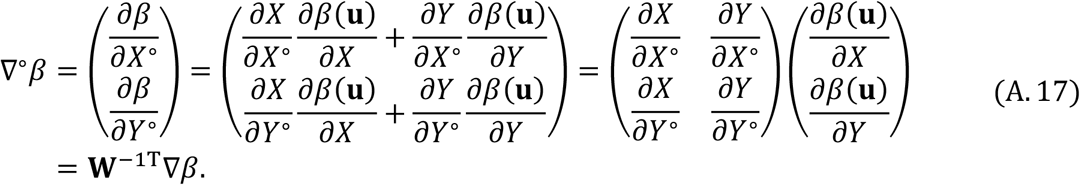

Substituting equation (A.15) with equation (A.17) into equation (A.14), we get

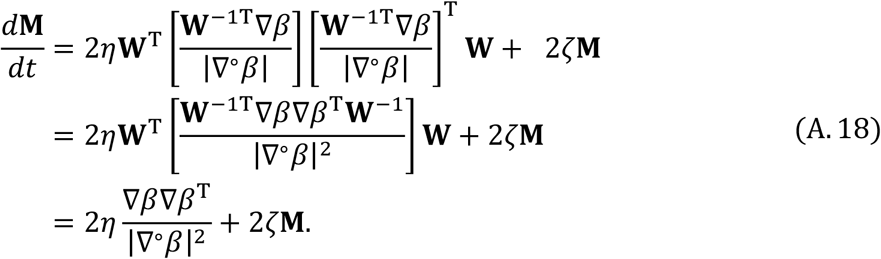

Now we assume that the unidirectional growth rate *η* is proportional to |∇^∘^*β*|^*k*^ with *k* being a positive integer:

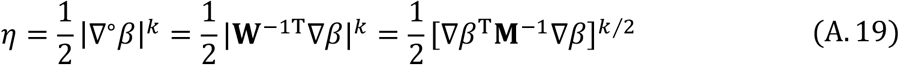

Substituting equation (A.19) into equation (A.18), we get

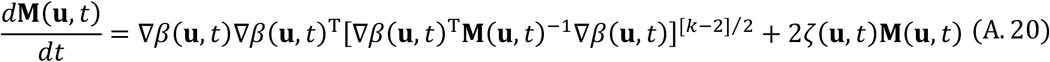

where the arguments “(**u**, *t*)” are recovered. Finally, assuming *k* = 2 and ζ(**u**, *t*) = 0 gives equation (3) in the main text. If ζ(**u**, *t*) and *β*(**u**, *t*) are constant along time, the growth metric at time τ is obtained as

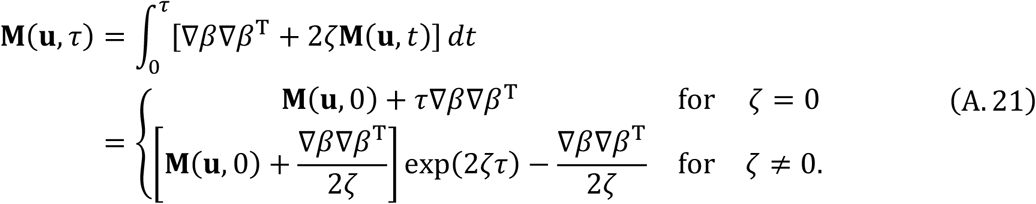

## Appendix B: Isometric embeddability

### B.1 Initial growth metric being identity

Here we show that the growth metric given by equation (6) in the main text,

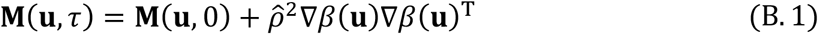

with 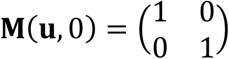, is identical to the growth metric calculated from the sheet shape given by equation (7),

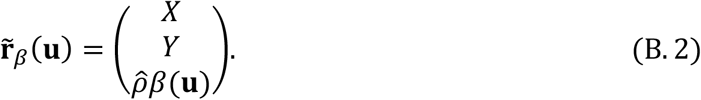

We confirm this relationship by deriving that the growth amount 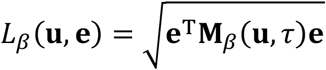 induced from equation (B.2) is identical to the growth amount 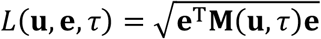 given by equation (B.1) for any **e**. Specifically, we see for **δu** = (*δX, δY*)^T^ and **e** = (*e*_*X*_ *e*_*Y*_)^*T*^ = **δu** / |**δu**| that

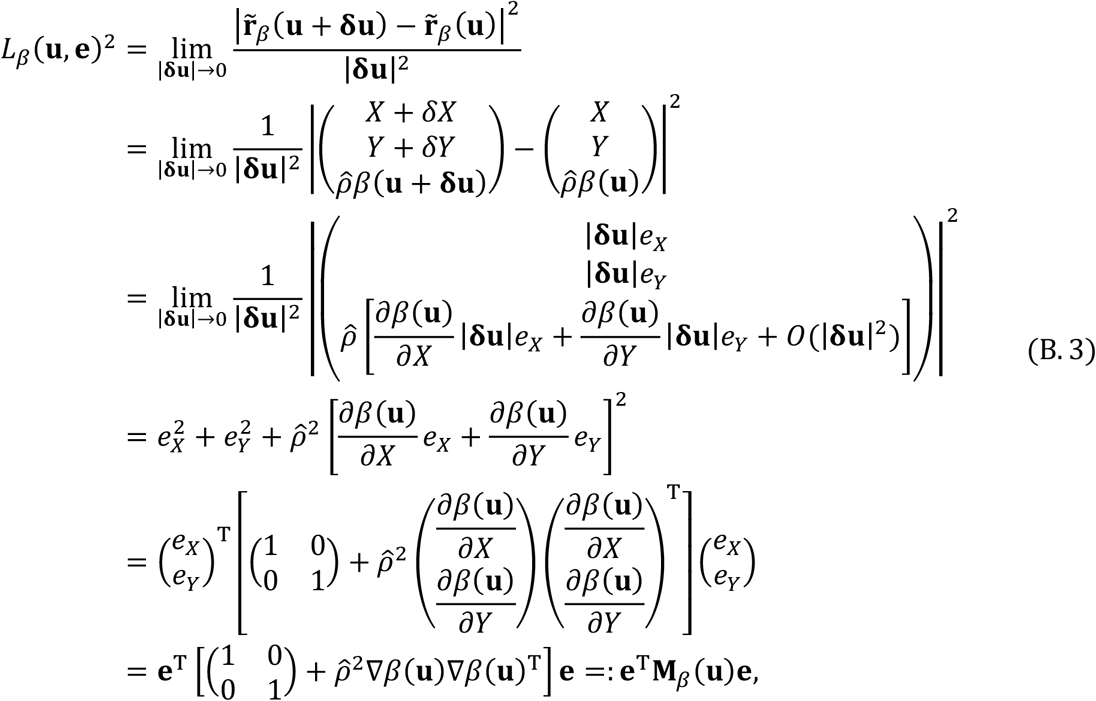

And which is identical to *L*(**u, e**, τ)^2^ = **e**^T^**M**(**u**, τ)**e** given by equation (B.1).

Therefore, as long as the basic shape for *β*(**u**) has a simple horn-like shape, we expect that the extended sheet calculated numerically using the finite element or vertex-spring methods, denoted by 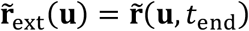, will have a shape similar to 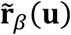, so that the realised growth amount 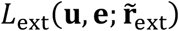 calculated from 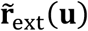 is similar to *L*_*β*_ (**u, e**). This means that the realised growth, denoted by the extension metric 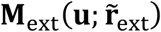, is similar to **M**_*β*_ (**u**), where 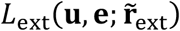 and 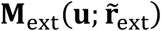 are defined as

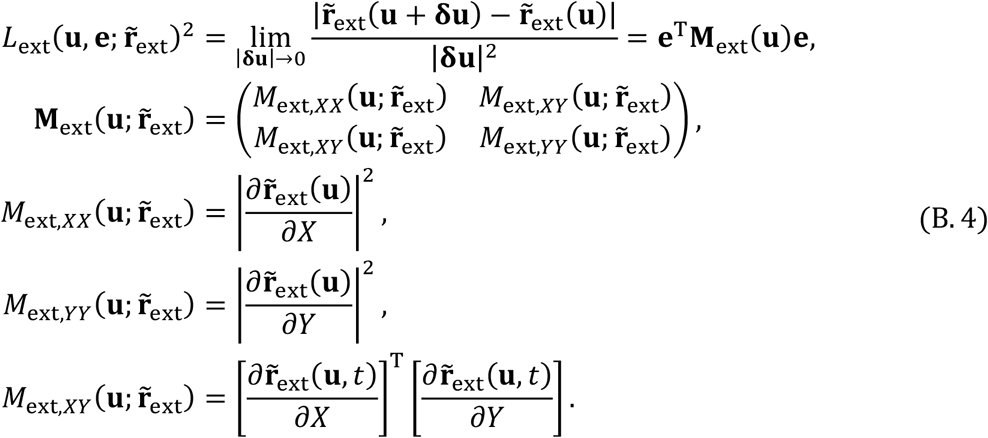

### B.2 Initial growth metric not being identity

Next, we consider the case that the initial growth metric **M**(**u**, 0) is not identity but constant over the sheet. Such a case arises when we fuse two basic shapes corresponding to *β*_1_(**u**) with a wide basic shape and *β*_2_(**u**) with a very narrow basic shape located at the middle of the slope of the basic shape of *β*_1_(**u**), as shown in figure 3. In this case, after the sheet growth induced by *β*_1_(**u**), the sheet around the basic shape of *β*_2_(**u**), denoted here by *β*(**u**), has approximately a constant growth metric, denoted here by **M**(**u**, 0).

An appropriate rotation of the material coordinates makes **M**(**u**, 0) diagonal with positive constant entries, which is expressed as

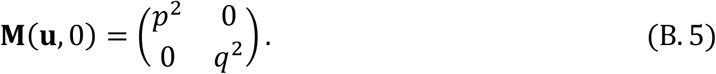

Then, as derived below, the growth metric 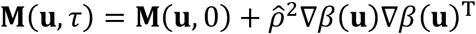 of the grown sheet has a smooth isometric embedding in the spatial coordinates given by

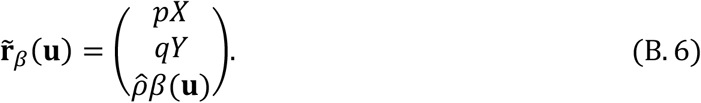

We confirm this in a manner analogous to the case of the initial growth metric being identity (Appendix B.1). Specifically, we see for **δu** = (*δX, δY*)^T^ and **e** = (*e*_*X*_ *e*_*Y*_)^T^ = **δu** / |**δu**| that

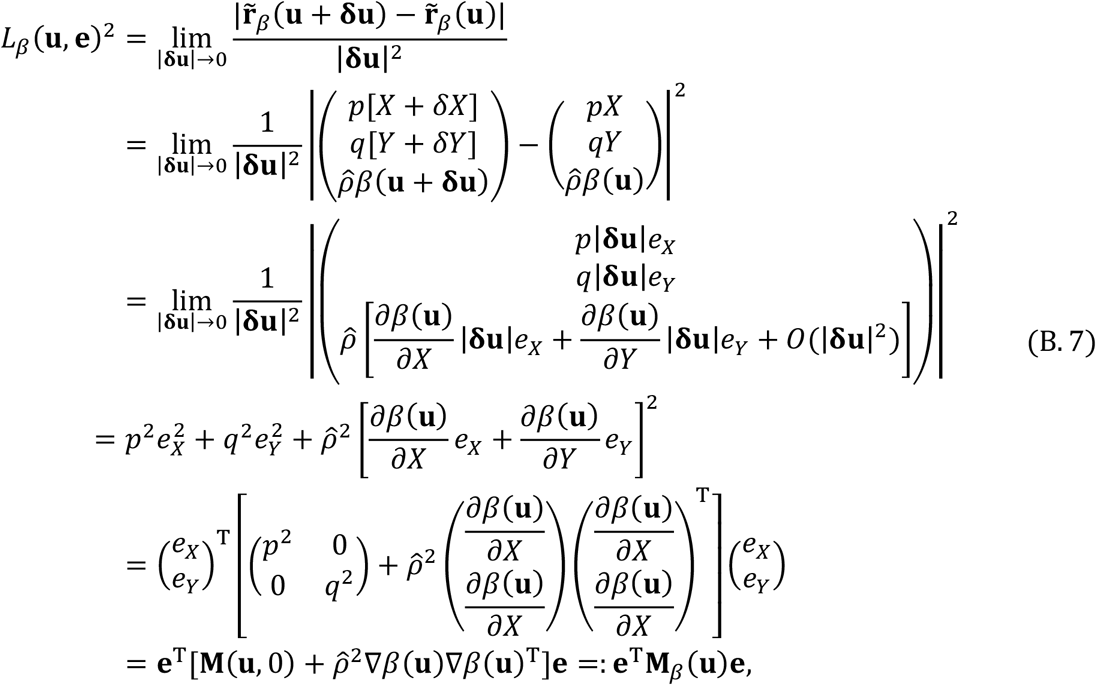

And which is identical to *L*(**u, e**, τ)^2^ = **e**^T^**M**(**u**, τ)**e**.

## Appendix C: Simulation method

### C.1 Finite element method

In the finite element method used in the present study, the epidermal sheet is divided into first-order hexagonal finite elements, and the spatial coordinates of vertices of these elements are all collected into a single vector **p**. The motion equation for **p** is written as

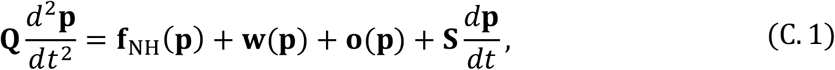

where **Q** and **S** respectively describe the mass and viscosity matrices. **f**_NH_ (**p**) is a vector describing the stress due to strain from the unstrained shape of each element, under the neo-Hookean hyper-elastic material model. **w**(**p**) is a vector describing water pressure from under the sheet. **o**(**p**) is a vector describing a uniform external force (parallel to the *z*-axis) acting on the edges of the sheet so that the entire sheet itself does not rise along with the *z*-axis even under the water pressure pushing up the sheet. This equation of motion is time-integrated with singly diagonally implicit Runge-Kutta (SDIRK) schemes, by using a C++ library MFEM-3.3.2.

In the initial state of the calculation, the finite elements were generated by equally dividing a square epidermal sheet along the *x* - and *y*-axes, where the sheet contains a single layer of elements along the *z*-axis. The time for the initial state is set at time *t* = τ (i.e., the time when the sheet growth ends), and the initial state for **p** corresponds to the 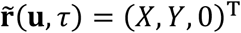. For each rectangular finite element, the unstrained lengths for the sides of the rectangles on the upper and lower surfaces are elongated according to the metric **M**_sim_(**u**, *t*). **M**_sim_(**u**, *t*) starts at time *t* = τ from its initial state **M**_init_(**u**) = **I** and then monotonically approaches the growth metric **M**(**u**) given by equation (6) or (9). After reaching **M**(**u**) at *t* = τ + *t*_ext_, **M**_sim_(**u**, *t*) is kept equal to **M**(**u**) until the calculation ends at *t* = τ + *t*_end_. Specifically, **M**_sim_(**u**, *t*) is defined as

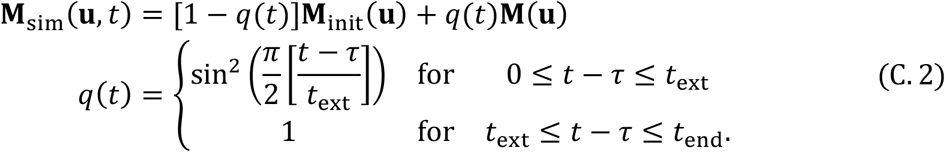

where the sine function simply represents that *q*(*t*) changes gently from zero to one in order to change the metric gradually. From **M**_sim_(**u**, *t*) the unstrained side length of an element connecting vertices *i* and *j* is given by

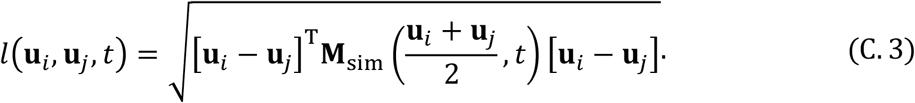

The strength of the water pressure at time *t* is given by *w*(*t*) = [1 − *q*(*t*)]*w*_init_ + *q*(*t*)*w*_end_ with constant parameters *w*_init_ and *w*_end_, where *w*_init_ is much larger than *w*_end_

[Parameters]

Initial sheet size: side length 8.0 along the *x*- and *y*-axes and thickness 0.2 along the *z*-axis

Time step: 0.5, Viscosity: 0.1

Bulk modulus: 5.0, Shear modulus: 0.05 (in Neo-Hookean hyperelastic model)

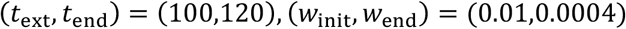

Fig. 2:

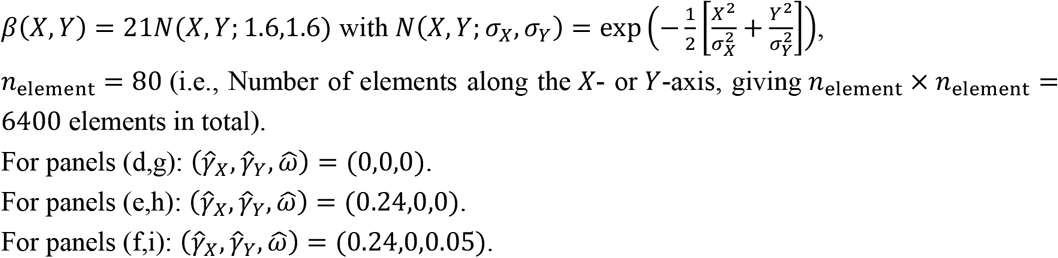

Fig. 3:

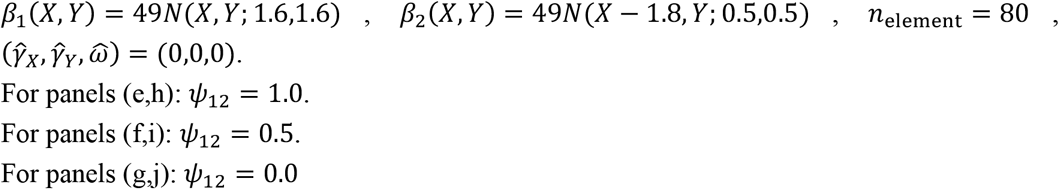

Fig. 4:

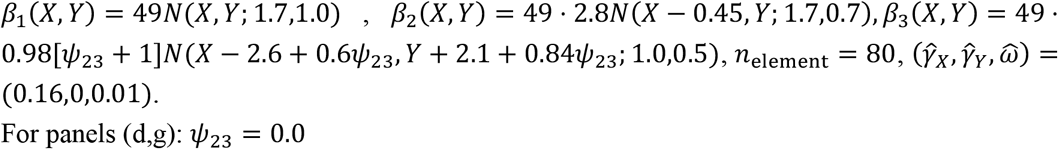

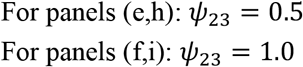

Fig. 5a,d,g:

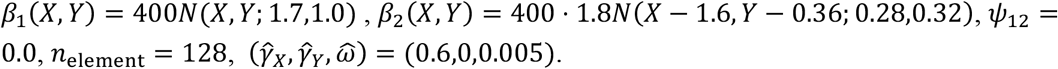

Fig. 5b,e,h:

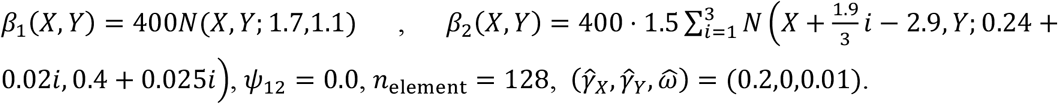

Fig. 5c,f,i:

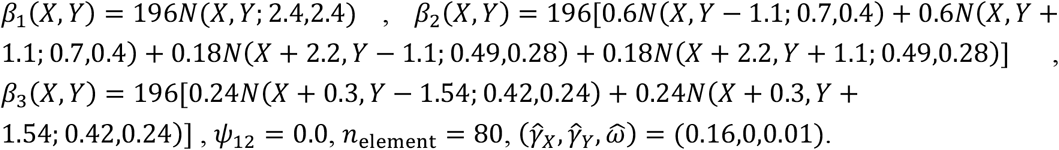

#### C.2 Vertex-spring method

[Main equation]

We assume that the epidermal sheet has no thickness and initially has a square shape with its sides being parallel with the *X*- or *Y*-axis in the material coordinates. We discretize the sheet into equally spaced vertices, denoted by (*i, j*) with *i, j* = 1, …, *N*, connected by springs. The spatial coordinates of vertex (*i, j*) are given by 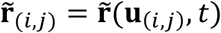. We describe the force acting on vertex (*i, j*) as

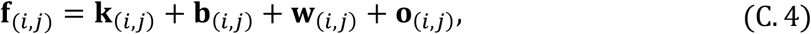

where **k**_(*i,j*)_, **b**_(*i,j*)_, **w**_(*i,j*)_, and **o**_(*i,j*)_ are forces due to springs (i.e., stretching elasticity), bending elasticity, the water pressure (from under of the sheet), and external force, respectively. We assume that the system is kept almost valanced so that |**f**_(*i,j*)_| is kept small at each time step, so that we can calculate the time evolution of the coordinates of vertex (*i, j*) by

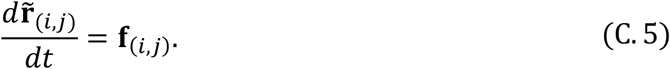

[Spring force, water pressure, and bending elasticity]

The spring force on vertex (*i, j*) is described as

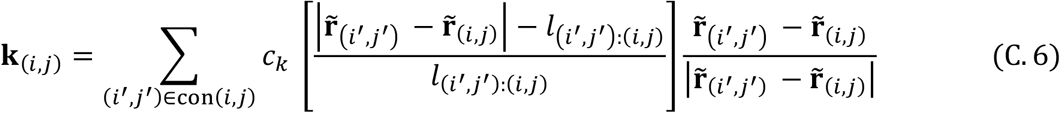

with a positive constant parameter *c*_*k*_ describing the spring constant per unit length, and where 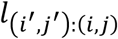 describes the spring length connecting vertices (*i, j*) and (*i*^′^, *j*^′^) chosen from the eight connected vertices, con(*i, j*) = {(*i* − 1, *j*), (*i* + 1, *j*), (*i, j* − 1), (*i, j* + 1)} ∪ {(*i* − 1, *j* − 1), (*i* + 1, *j* − 1), (*i* − 1, *j* + 1), (*i* + 1, *j* + 1)}. 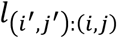 at time *t* is given by

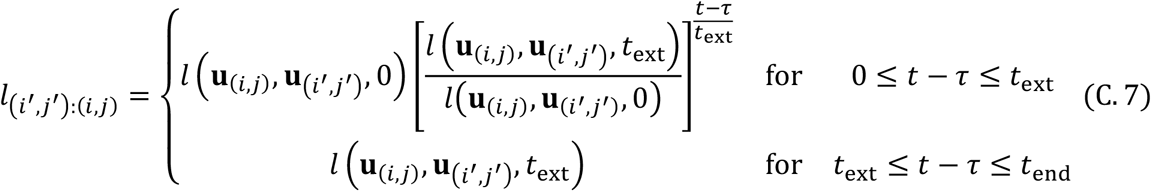

with 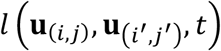 defined in in equation (C.3) with *t* = *τ* + *t*_ext_

The bending-elastic force **b**_(*i,j*)_ on the vertex (*i, j*) is described as the sum of the bending-elastic force 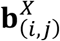 along the *X*-axis: {(*i* − 1, *j*), (*i, j*), (*i* + 1, *j*)} and the bending-elastic force 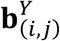 along the *Y*-axis: {(*i, j* − 1), (*i, j*), (*i, j* + 1)}. Specifically, we describe **b**_(*i,j*)_ as:

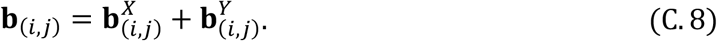

We denote by 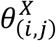 the bending angle of the sheet at vertex (*i, j*) along *X*, as illustrated in figure 9a. The 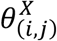 is the angle between the segment (*i* − 1, *j*): (*i, j*) and the segment (*i, j*): (*i* + 1, *j*), given by the inner product of the corresponding two unit vectors:

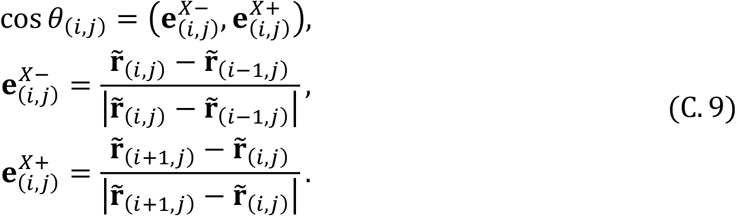

**Figure 9.**
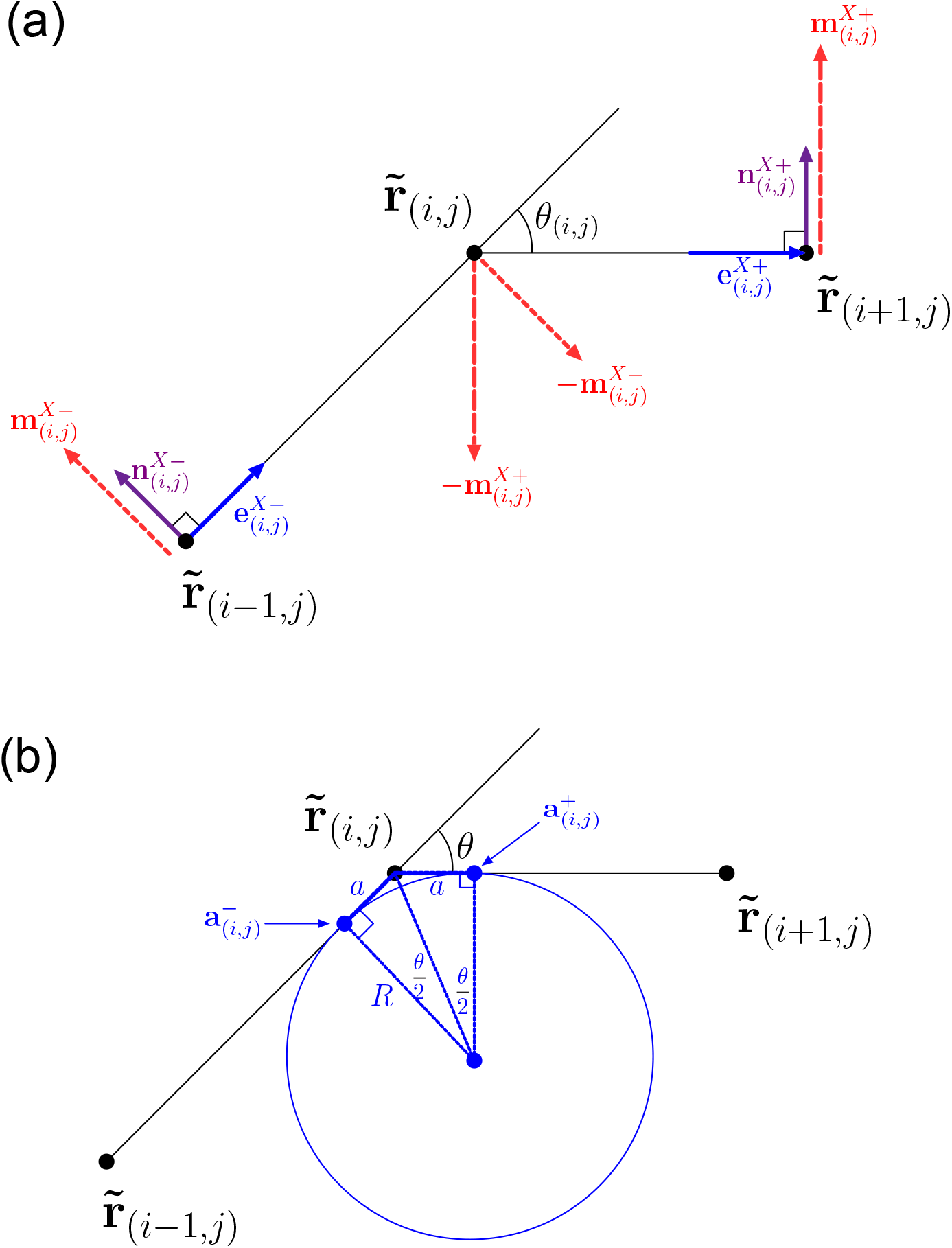
Bending elasticity of a sheet modeled with the vertex-spring method. (a) Illustration of action and reaction forces due to bending elasticity of the sheet. (b) Degree of bending described with curvature radius.

The 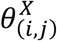 produces the forces 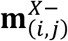 and 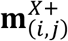 on vertices (*i* − 1, *j*) and (*i* + 1, *j*), respectively. At the same time, due to the law of action and reaction, vertex (*i, j*) undergoes 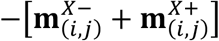. We assume that the bending elasticity caused by the bending angle *θ*_(*i,j*)_ is proportional to the inverse of the curvature radius 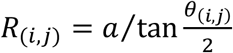 (illustrated in figure 9b, where the corresponding circle is tangent to line segments (*i* − 1, *j*): (*i, j*) and (*i, j*): (*i* + 1, *j*) respectively at points 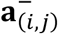 and 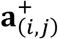 both of which have the constant distance *a* from 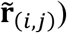. To keep 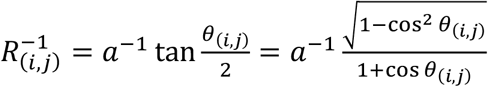 from being infinite, we introduce a constant saturation parameter *c*_b*s*_ > 0 and describe the torque as

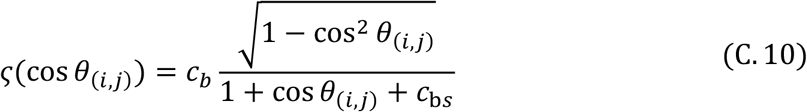

where *a*^−1^ is included in a positive constant parameter *c*_b_. Then 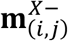 and 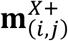 are calculated as

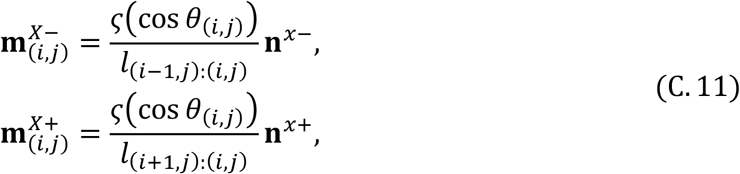

where 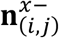 and 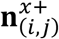 are unit vectors included in the plane that contains the three vertices 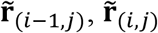, and 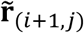 and are orthogonal to 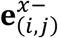 and 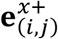, respectively. Specifically, 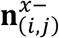 and 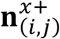 are given by

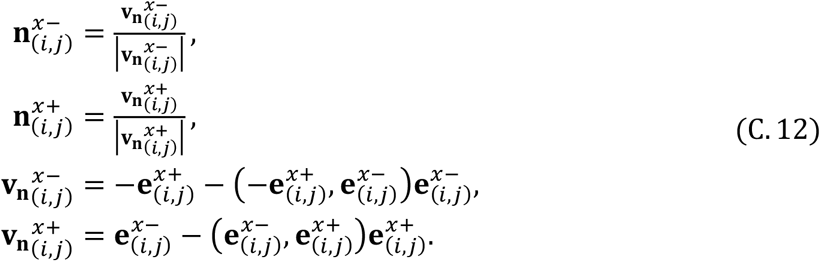

On this basis, we obtain the bending-elastic force 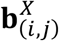 on vertex (*i, j*) along the *X*-axis as the sum of 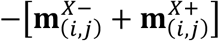 from the bending at vertex (*i, j*), 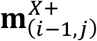 from (*i* − 1, *j*), and 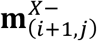 from (*i* + 1, *j*):

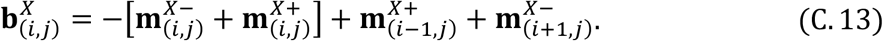

Analogously, we obtain the bending-elastic force 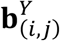 on vertex (*i, j*) along the *Y*-axis from equations (C.9-C.13) with replacement of the superscript “*X*” with “*Y*.” Then, the total bending elastic force **b**_(*i,j*)_ on vertex (*i, j*) is calculated by equation (C. 8).

The force exerted by water pressure on the vertex (*i, j*) is described as

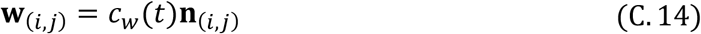

with a positive time-dependent parameter *c*_W_(*t*) defined with a sigmoid function,

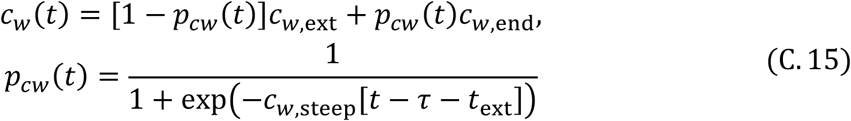

with constant parameters *c*_W,ext_, *c*_W,end_, and *c*_W,steep_. **n**_(*i,j*)_ is the normal vector of the sheet at 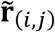, given by the average of the four normal vectors of the four triangles sharing vertex (*i, j*) (i.e., {(*i, j*), (*i* + 1, *j*), (*i, j* + 1)}, {(*i, j*), (*i, j* + 1), (*i* − 1, *j*)}, {(*i, j*), (*i* − 1, *j*), (*i* − 1, *j* − 1)}, and {(*i, j*), (*i* − 1, *j*), (*i* + 1, *j* − 1)}.

Finally, the upward external force on the vertices (*i, j*) (acting only on the sheet edges) could be described as follows:

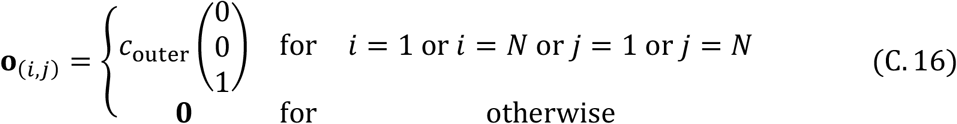

with a positive constant parameter *c*_outer_.

[Parameters]

Sheet size: 4.0 (along *X*-axis), 4.0 (*Y*-axis), 0.0 (*Z*-axis)

Number of vertices: 129×129 (along *X*- and *Y*-axes)

Time step: Δ8.0 × 10^− 4^ (Explicit Euler method)

Elasticity parameters: *c*_l_ = 10, *c*_b_ = 0.23, *c*_bs_ = 0.0001, *c*_outer_ = 40

Morphogen distribution: 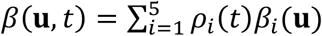 with *ψ*_*ij*_ = ∫ *ρ*_i_(*t*)*ρ*_*j*_(*t*)*dt* = 0 for all *i* ≠ *j*

Function for linear combination of similar Normal distributions:

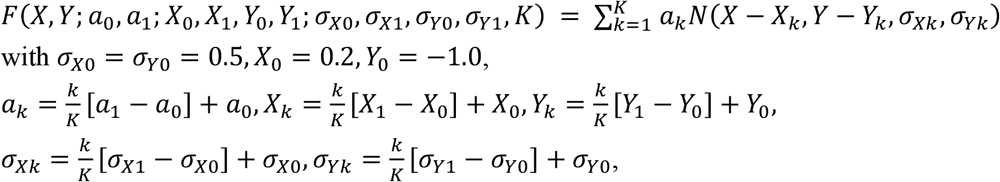

with *σ*_*X*0_ = *σ*_*Y*0_ = 0.5, *X*_0_ = 0.2, *Y*_0_ = −1.0,

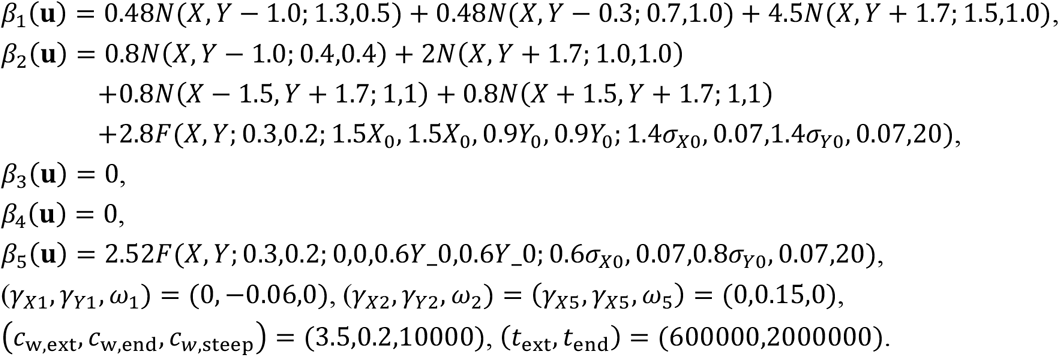

Fig. 6:

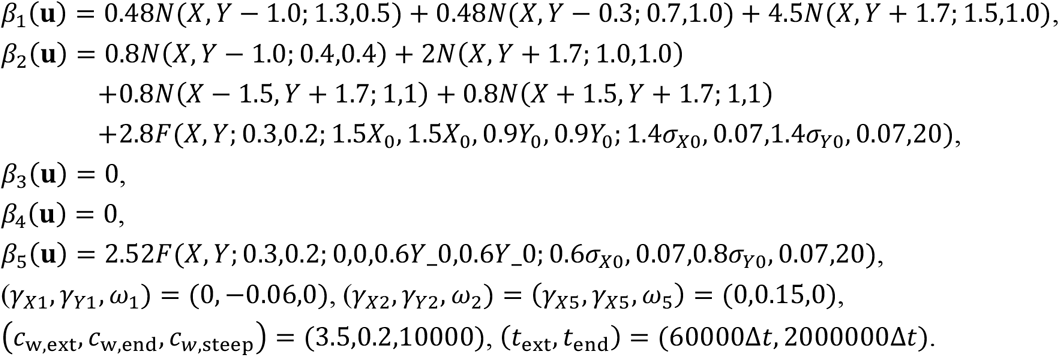

Fig. 7:

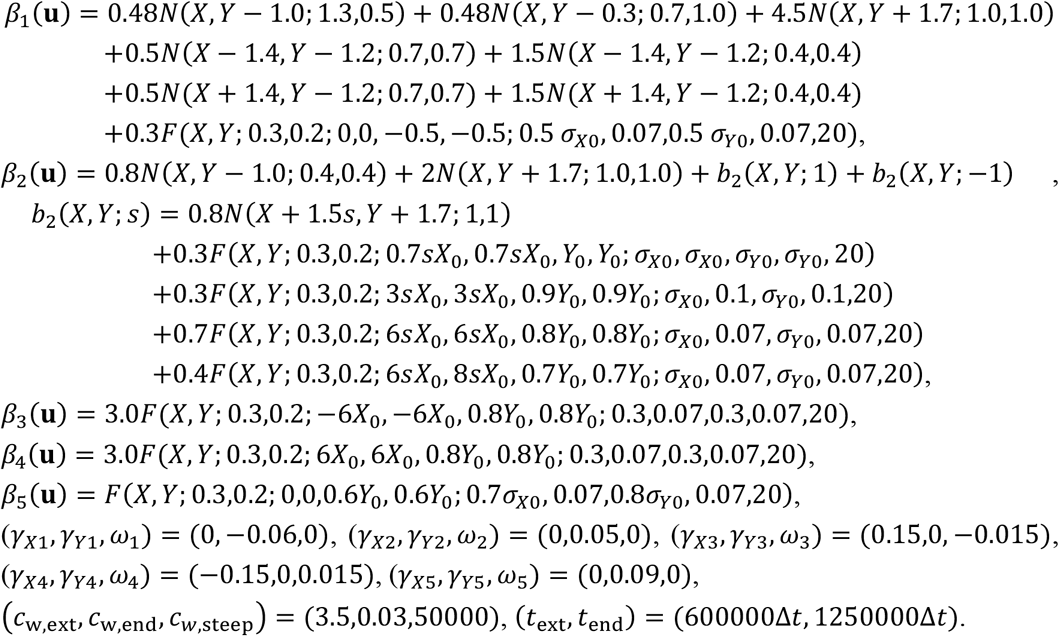

Fig. 8:

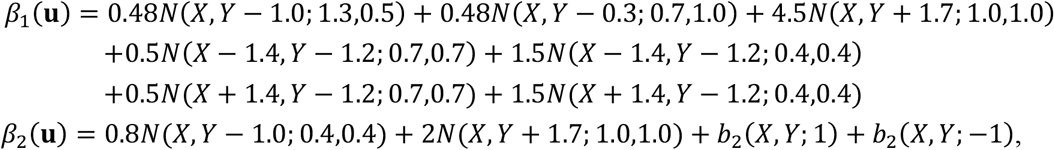

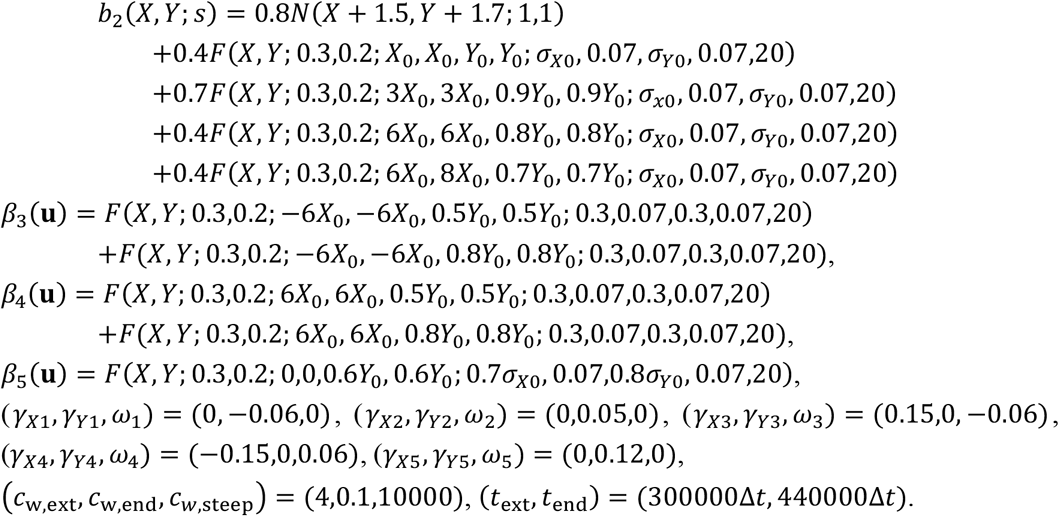

## Appendix D: Modification of extended shapes

We can curve and twist the shape of the extended sheet corresponding to the basic shape for morphogen distribution *β*(**u**), by introducing two additional kinds of morphogens, denoted by *γ* and ω. These additional morphogens can modify the sheet growth regulated by *β*(**u**), as explained below.

For simplicity of explanation, we assume that *β*(**u**) has a simple horn-like basic shape and has a peak at **u** = (*X,Y*)^T^ = (0,0)^T^ in the material coordinates, so that it is symmetric with respect to both the *X*- and *Y*-axes (figure 2a). If the sheet elements in the region of *X* < 0 grow more than those in *X* > 0 (Fig. 2e), this horn-like shape can be curved in the positive *X*-direction. As the morphogen gradient in the *X*-direction, ∂*β*(**u**)/∂*X*, is positive for *X* < 0 and negative for *X* > 0, the curved horn can also be attained if the region of ∂*β*(**u**)/∂*X* > 0 grows more than that of ∂*β*(**u**)/∂*X* < 0. Specifically, we get the curved horn by modifying equation (3) into

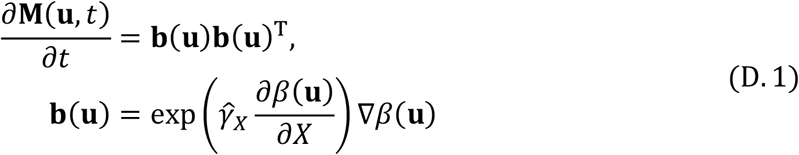

with a constant 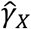 describing the direction and strength of the curve. Note that equation (D.1) applies even when the peak of the basic shape is not located at the origin. We can curve the horn-like shape along a direction other than the *X*-axis, by introducing the morphogen distribution *γ*(**u**) with its gradient being constant, given by 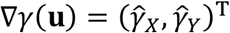, and by extending **b**(**u**) into

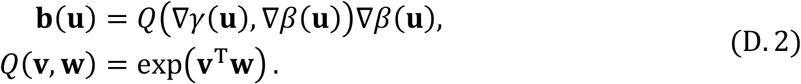

Regarding the function *Q*(**v, w**), a function other than E*x*p(**v, w**) may also be used, as long as it is no less than 1 and monotonically increases with respect to |**v**| and 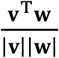. In our present study, *Q*(**v, w**) = E*x*p(**v**^T^**w**) was used for figures 1-5, and 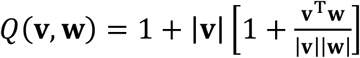 was used for figures 6-8.

The curved horn-like shape attained by equation (D.1) or (D.2) can be further twisted by slight rotation of the local growth direction at each position of the sheet. We can achieve this by introducing a uniform morphogen distribution 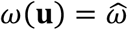 with a constant 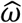, and by further modifying equation (D.2) into

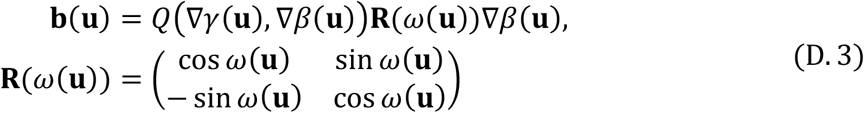

(figure 2f,i).

For fusion of the two basic shapes corresponding to *β*_1_(**u**) and *β*_2_(**u**) (corresponding to equation (8) in the main text), we assume that these basic shapes can have different degrees of curves (given by *γ*_1_(**u**) and *γ*_2_(**u**)) and twists (given by ω_1_(**u**) and ω_2_(**u**)). Then, we define **b**(**u**, *t*) by

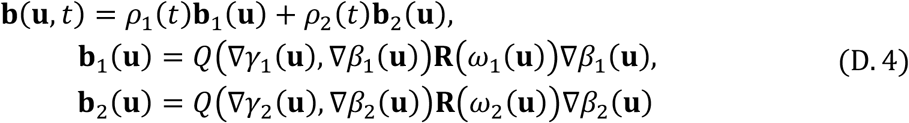

with 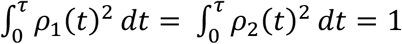. In this case, we see that

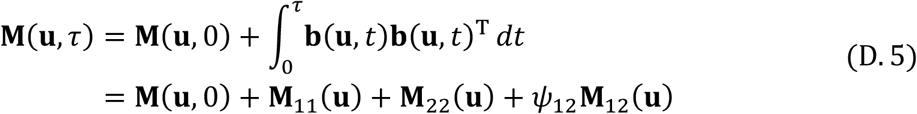

with

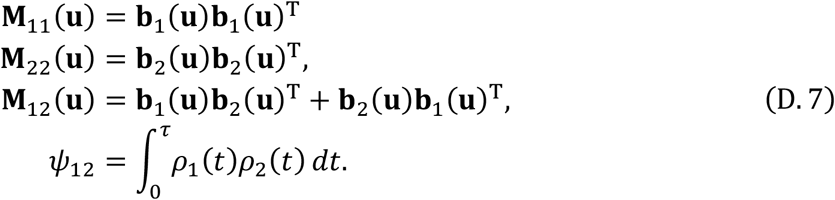

In figure 2 in the main text, 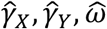 are chosen as 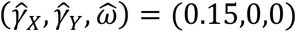 for (e) and (h), and 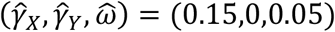 for (f) and (i).

